# Regulatable assembly of synthetic microtubule architectures using engineered MAP-IDR condensates

**DOI:** 10.1101/2023.03.14.532644

**Authors:** Chih-Chia Chang, Scott M. Coyle

**Affiliations:** Biophysics Graduate Program University of Wisconsin-Madison, Madison, WI, 53705, USA; Department of Biochemistry University of Wisconsin-Madison, Madison, WI, 53705, USA

## Abstract

Microtubules filaments are assembled into higher-order structures and machines critical for cellular processes using microtubule-associated proteins (MAPs). However, the design of synthetic MAPs that direct the formation of new structures in cells is challenging, as nanoscale biochemical activities must be organized across micron length-scales. Here we develop synthetic MAP-IDR condensates (synMAPs) that provide tunable and regulatable assembly of higher-order microtubule structures *in vitro* and in mammalian cells. synMAPs harness a small microtubule-binding domain from oligodendrocytes (TPPP) whose activity can be synthetically rewired by interaction with condensate-forming IDR sequences. This combination allows synMAPs to self-organize multivalent structures that bind and bridge microtubules into synthetic architectures. Regulating the connection between the microtubule-binding and condensate-forming components allows synMAPs to act as nodes in more complex cytoskeletal circuits in which the formation and dynamics of the microtubule structure can be controlled by small molecules or cell-signaling inputs. By systematically testing a panel of synMAP circuit designs, we define a two-level control scheme for dynamic assembly of microtubule architectures at the nanoscale (via microtubule-binding) and microscale (via condensate formation). synMAPs provide a compact and rationally engineerable starting point for the design of more complex microtubule architectures and cellular machines.

## Introduction

Microtubules (MTs) are one of the major filament systems of the eukaryotic cytoskeleton, extending throughout the cell to build up internal structure, position organelles at specific locations, and facilitate vesicle transport. Individual MTs are further organized into higher-order structures and machines critical for cellular processes, such as the mitotic spindle or ciliary axonemes^1–4^. The assembly and function of these structures depends on the coordinated action of microtubule-associated proteins (MAPs), proteins that bind MTs and work across cellular length scales to direct the organization of a specific target structure^5,6^.

The diversity of microtubule-based structures and functions seen across cells in nature suggests that synthetic MAPs could be engineered to assemble new microtubule structures with novel functions inside cells, analogous to how synthetic signaling proteins can rewire cellular information processing^7–10^. However, because MTs are the largest filaments of the eukaryotic cytoskeleton, with a diameter of 25 nm and more than 1600 tubulin subunits per micron^11–13^, it is not obvious how to best engineer a synthetic MAP protein such that its nanoscale activities can be effectively integrated across the microscale within cells.

Protein condensation mediated by phase separation has emerged as a mechanism by which nanoscale components can be organized and condensed into micron-scale droplets to drive biochemical interactions and chemistry in cells^14–17^. Although these droplets bear little resemblance to the ordered structures of the cytoskeleton, recent studies suggest roles for condensates in amplifying the activity MAPs associated with microtubule bundling, branching, and tip growth such as tau, TPX2, NuMA, and +TIPs^18–21^. These observations are intriguing, as the high concentrations and reduced-order of condensates could provide the multimerization, valency and flexibility needed to bridge length-scales for synthetic applications. However, these specific MAPs are poor candidates for synthetic engineering, as they are large multi-domain proteins that work in multi-component networks to regulate cell-essential structures such as the mitotic spindle^22–24^**(Fig. 1A)**.

**Fig. 1.**
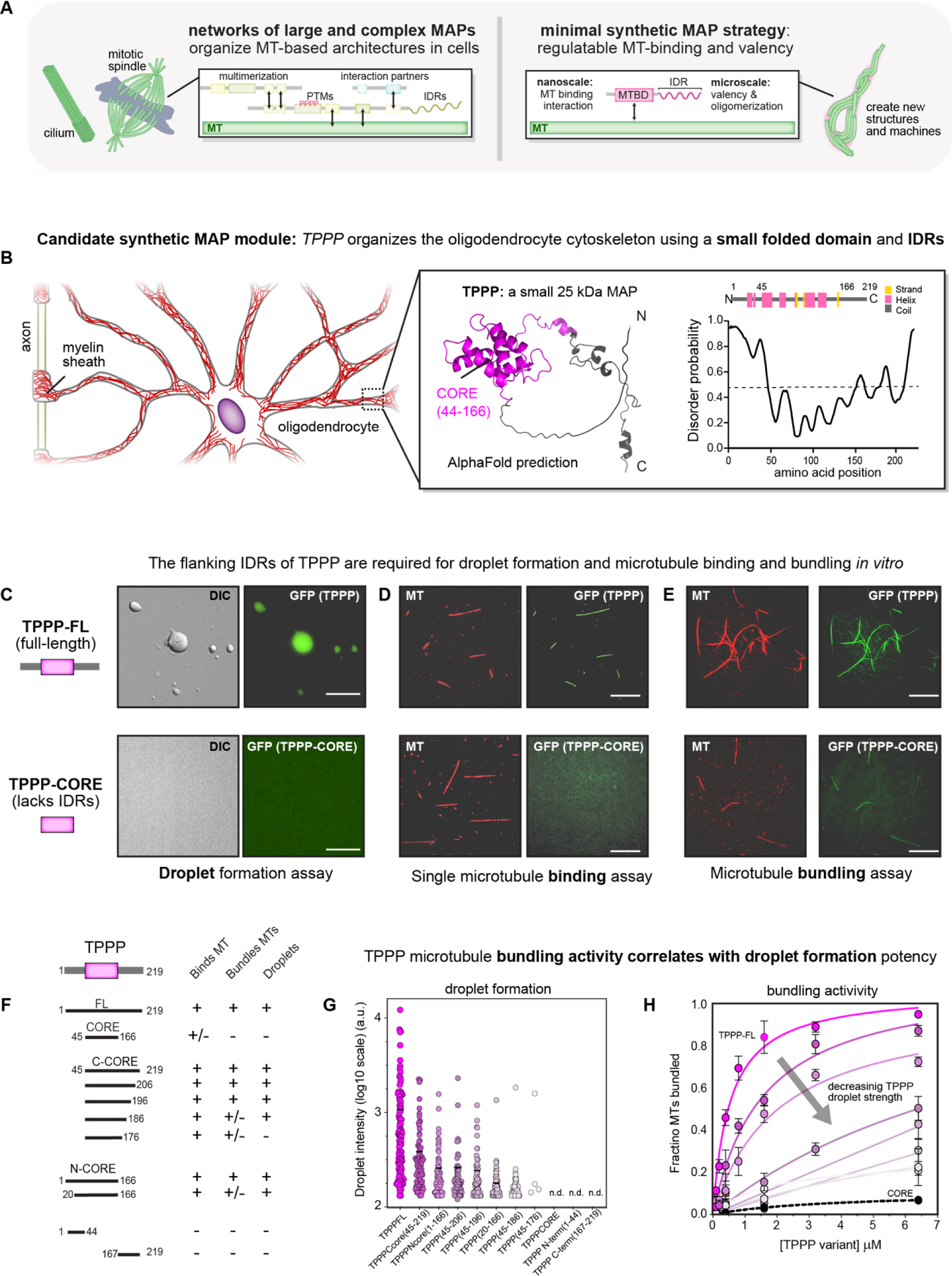
Identification of TPPP as a condensate-regulatable microtubule bundling module for building synthetic microtubule assemblies. (A) Overview for designing synthetic microtubule-associated proteins (MAPs) to build new microtubule architectures. (left) Native MAPs, such as spindle assembly factors (e.g., NuMA and TPX2), have multi-domain architectures with many activities that provide valency, multimerization, and complex dynamic regulation; (right) a minimal design for a regulatable synMAP protein based on a single microtubule-binding domain to facilitate microtubule attachment and a condensate-forming sequence (IDR) to drive valency and multimerization. (B) Identification of TPPP as a candidate module for synthetic engineering using the strategy in (A). TPPP is a 24 kDa protein that promotes microtubule nucleation, bundling, and stabilization in oligodendrocytes. A 3D visualization of the AlphaFold structure prediction for TPPP is shown, with the folded CORE domain in pink and unstructured regions in gray. The corresponding secondary structure prediction of TPPP protein domains is indicated, with α-helical and β-strands structures are shown as pink and yellow blocks, along with the PrDOS-predicted intrinsic disorder of TPPP. Higher scores indicate a greater degree of disorder. (C) Epifluorescence images of TPPP droplet formation. TPPP, but not the isolated CORE domain, phase-separate into drops in the presence of 12% of dextran crowding agent. DIC (left) and green fluorescence (right) images of GFP-TPPP FL and GFP-TPPP-CORE, both at the concentration of 20 μM with 20 mM HEPES, 50 mM NaCl, 3 mM DTT, and 12% dextran. Scale bars, 10 μm. (D) TIRF images of single-microtubule binding for TPPP and isolated CORE domain. GMPCPP-stabilized microtubules (Alexa-594 and biotin-labeled) immobilized on a glass surface, were incubated with GFP-TPPP FL and GFP-TPPP-CORE, respectively. Both at the concentration of 600 nM. Scale bars, 10 μm. (E) TIRF microscopy images of microtubule bundling for TPPP and isolated CORE domain. GFP-TPPP FL and GFP-TPPP-CORE (green) were mixed with in vitro polymerized Alexa-594 (red), biotin-labeled, and GMPCPP-stabilized microtubules for 5 min, then captured microtubule bundles by NeutrAvidin-coated biotin coverslips. Both at the concentration of 6.4 μM. Scale bars, 10 μm. (F) Behavior of different TPPP truncations and fragments with respect to MT-binding, bundling, and droplet formation activities. Symbols denote experiments performed qualitatively. (G) Quantification of GFP-TPPP variants from (F) with respect to droplet fluorescence signal intensity. The mean was determined from data pooled from five representative images. GFP-TPPP variants droplet number in total five images, respectively; TPPP FL, n = 107; TPPP - Ccore(45-219), n = 92; TPPP-Ncore(1-166), n = 73; TPPP(45-206), n = 53; TPPP(45-196), n = 60; TPPP(20-166), n = 89; TPPP(45-186), n = 90; TPPP(45-176), n = 4. TPPP-CORE(45-166), TPPP N-term, and TPPP C-term showed n.d., not detectable droplets. All GFP-TPPP variants were in the concentration of 20 μM with 20mM HEPES, 50mM NaCl, 3mM DTT, and 12% dextran. (H) MT bundling activity for the variants in (F) was quantified as an EC50 from full titration curves measuring the fraction of microtubules in bundles as a function of variant concentration. TPPP FL, EC50 = 0.54 μM; TPPP-Ccore, EC50 = 1.58 μM; TPPP-Ncore, EC50 = 2.06 μM; TPPP(45-206), EC50 = 10.58 μM; TPPP(45-196), TPPP(45-186), TPPP(20-166), and TPPP-CORE curves and EC50 constants were determined by fitting to a hyperbola. Mean (points) and SEM (error bars) of three replicate experiments (error bars) shown.

We sought a non-essential condensate-regulatable MAP with a simplified domain architecture that could provide a starting point assembling synthetic MT architectures. Tubulin polymerization-promoting protein (TPPP) is a small 24 kDa MAP found in oligodendrocytes that contains only a single folded “CORE” domain^25,26^ **(Fig. 1B)**. Despite its small size, TPPP is proposed to nucleate and stabilize MTs at discrete structures called Golgi outposts to promote elongation, branching, and polarization of myelin sheaths in oligodendrocytes^27^. Pathologically, TPPP aggregates are associated with Parkinson’s disease and Multiple System Atrophy, and reversing TPPP aggregation has been proposed as a therapeutic strategy^28–30^. Critically, the structured domain of TPPP is flanked by low-complexity intrinsically disordered sequences (IDRs), suggesting a possible role for protein condensation in its activity.

Here we take advantage of TPPP’s minimal architecture and desirable features to develop synMAPs, engineered proteins that provide tunable and regulatable assembly of higher-order microtubule structures in living cells **(Fig. 1A)**. Our designs are based on our observation that TPPP’s native MT binding and bundling activity is effectively tuned by its intrinsic ability to form protein condensates. Both *in vitro* and in mammalian cells, we show that microtubule bundling activity can be synthetically re-wired by fusing TPPP fragments to other unrelated condensate-forming IDR sequences. This basic control logic can be exploited to engineer more elaborate circuits that regulate the connection between TPPP and IDRs, allowing large MT architectures to be assembled and controlled in response to small molecules and cell-signaling inputs. By systematically testing a panel of synMAP circuits that vary the strength of the microtubule-binding domain or the IDR, we experimentally define a two-level design manual for controlling synMAP-directed microtubule assembly at the nanoscale and microscale. synMAPs offer a simple and easily engineered module for designing synthetic microtubule architectures and machines *in vitro* and inside living cells.

## Results

### Identification of TPPP as a condensate-regulatable microtubule bundling module for building synthetic microtubule assemblies

Tubulin polymerization-promoting protein (TPPP) is a small 24 kDa MAP found in oligodendrocytes that has been shown to bind and bundle microtubules (MTs) *in vitro*. Inspection of TPPP’s amino acid sequence and AlphaFold structure prediction revealed a folded ’’CORE’’ domain flanked by unstructured N- and C-terminal regions (IDRs) **(Fig 1B)**. As IDRs are common features of phase separating proteins, this suggested that TPPP might represent a compact condensate-regulatable starting point for engineering synthetic MAPs. To determine the relationship between TPPP’s IDRs and its biological activity, we applied three *in vitro* assays in parallel: 1) a droplet formation assay to determine whether a TPPP construct could phase separate in the presence of crowding reagents; 2) a TIRF-based assay for measuring binding to immobilized single MTs in the absence of crowders; and 3) a TIRF-based bundling assay that measures the fraction of microtubules in bundles as a function of TPPP concentration to define an EC50 for quantitative comparison (detailed in methods)^18,22,31^.

We first expressed and purified GFP-tagged full-length TPPP and the isolated TPPP-CORE domain (aa.45-166) which completely lacks the unstructured N and C termini─and tested each construct for droplet formation. We found that TPPP-FL phase-separated to form drops under a wide-range of different molecular crowding conditions **(Fig. 1C).** In FRAP experiments, TPPP-FL droplets showed clear fluorescent recovery over time, suggesting liquid-like properties **(Supplementary Fig. b,c and Supplementary Video S1)**. In contrast, no droplets were observed for TPPP-CORE under any conditions tested. Nearly identical results were observed using untagged constructs and DIC microscopy to visualize these droplets **(Fig. 1C and Supplementary Fig. 1a)**. TPPP thus forms protein droplets *in vitro* that depend on the presence of its flanking IDRs.

We then tested if TPPP droplet formation correlated with microtubule binding or bundling activity, measured in the absence of any molecular crowders. Droplet-competent TPPP-FL strongly decorated single-MTs immobilized on coverslips at concentrations as low as 600 nM, while the isolated CORE domain showed weak binding along MTs only at concentrations greater than 6 μM **(Fig. 1D and Supplementary Fig 2).** Similarly, while TPPP-FL showed strong MT bundling activity (EC_50_=0.54 μM), TPPP-CORE bundling was barely detectable even at concentrations greater than 6 μM **(Fig. 1E)**. Thus, TPPP’s *in vitro* MT binding and bundling activity correlates with its ability to form droplets.

We next applied these assays to a series of TPPP truncations to quantitatively characterize the relationship between condensate formation and TPPP activity. Neither the N-terminal IDR (aa.1-44) nor the C-terminal IDR (aa.167-219) sequences were sufficient on their own to form droplets, bind, or bundle MTs **(Fig. 1F,G and Supplementary Fig 2b).** However, when fused to the TPPP-CORE domain, both the N-terminal IDR (TPPP N-CORE) or the C-terminal IDR (TPPP C-CORE) could support droplet formation, MT binding, and MT bundling. Fine-scale truncations of the N-terminal IDR and C-terminal IDRs in this context showed that droplet size scaled with the length of the IDR extension **(Fig. 1G, H)** and correlated with performance in our MT-bundling and single-MT binding assays (**Fig. 1F-H and Supplementary Fig 2b)**. Together these data suggest that TPPP possesses a condensate-regulatable activity that can be quantitatively tuned by the magnitude of droplet formation.

### TPPP bundling activity can be rewired by fusion to orthogonal condensate-forming sequences to construct synMAP proteins

Because the isolated TPPP-CORE domain has only weak MT-binding activity **(Fig. 1E and Supplementary Fig 2b)**, we reasoned that a function for condensates in this system could be to create the multimerization and high-valency needed to bridge multiple weak binding events across multiple MTs into bundles. If true, it should be possible to engineer synthetic MAPs (synMAPs) that replace TPPP’s endogenous IDRs with alternative droplet-forming sequences unrelated to MT biology. To test, we designed and characterized synMAP TPPP variants in which we fused TPPP-CORE to well-characterized phase-separating IDRs: the IDRs of DDX4 (aa.1-236), FUS (aa.1-214), and LAF-1 RGG (aa.1-200)^32–36^. These synMAP TPPP chimeras provide a control knob for probing how condensate formation can be harnessed to control and tune MT bundling activity **(Fig. 2A)**.

**Fig. 2.**
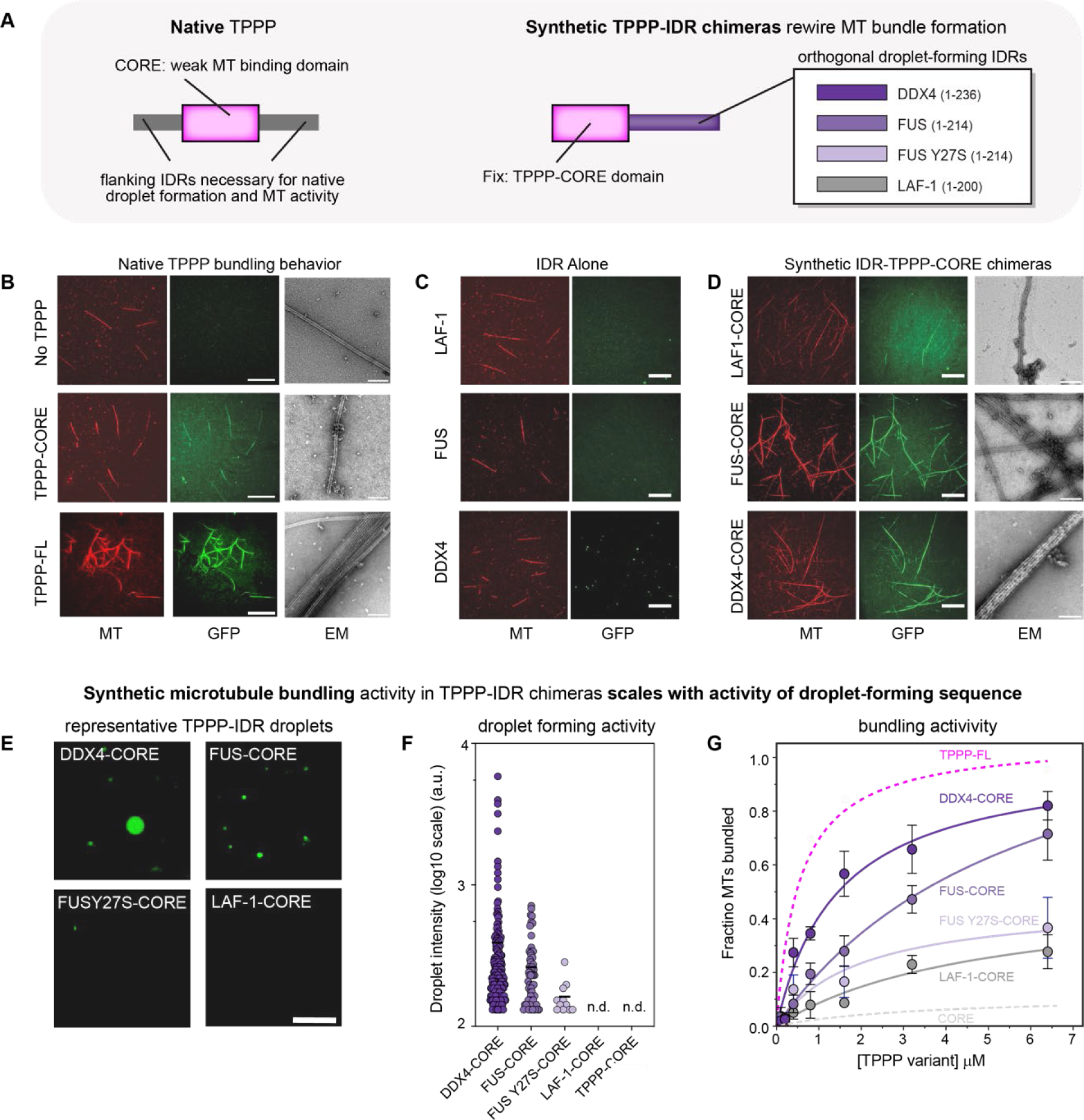
TPPP bundling activity can be rewired by fusion to orthogonal condensate-forming sequences to construct synMAP proteins. (A) Strategy for rewiring TPPP activity using orthogonal IDR sequences to construct synMAPs. TPPP structured CORE domain (left) can act as a weak MT-binding domain but bundle formation requires the presence of one of flanking intrinsically disordered regions (IDRs). Schematic of synMAP design (right): the IDRs of DDX4, FUS, FUSY27S (mutant), and LAF-1 were fused with TPPP-CORE domain as shown in table. These TPPP-IDR chimeras were tagged with GFP at their N-terminus. (B) TIRF and EM images showing the *in vitro* bundling behavior of native TPPP-FL and the isolated TPPP-CORE domain. Free microtubules are shown as a negative control. Scale bars are 20 μm (TIRF images) and 200 nm (EM images). (C) TIRF images showing that condensate-forming sequences (IDRs of DDX4, FUS, and LAF-1) do not interact with or bundle microtubules *in vitro*. Scale bars, 20 μm. (D) TIRF and EM images showing the *in vitro* bundling behavior of synMAP TPPP-IDR chimeras. Scale bars are 20 μm (TIRF images) and 200 nm (EM images). (E) Representative fluorescent images of droplets formed by synMAP TPPP-IDR chimeras *in vitro*. Assays were performed at a protein concentration of 20 μM with 20 mM HEPES, 50 mM KCl, 3 mM DTT, and 12% dextran. Scale bars, 10 μm. (F) Quantification of synMAP TPPP-IDR chimeras droplet fluorescence signal intensity. The mean was determined from data pooled from five different images. Droplet number was respectively: DDX4-CORE, n = 179; FUS-CORE, n = 57; FUSY27S-CORE, n = 11; LAF-1-CORE, n = 0; TPPP-CORE, n = 0. All synMAP TPPP-IDR chimeras were tested at the same concentration. (G) MT bundling activity for the synMAP TPPP-IDR chimeras in (F) was quantified as an EC50 from full titration curves measuring the fraction of microtubules in bundles as a function of variant concentration. TPPP FL, EC50 = 0.54 μM, data was added as a point of comparison. DDX4-CORE, EC50 = 1.47 μM; FUS-CORE, EC50 = 5.92 μM; FUSY27S-CORE, and LAF-1-CORE curves and EC50 constants were determined by fitting to a hyperbola. Mean (points) and SEM (error bars) of three replicate experiments shown.

On their own, the IDRs from DDX4, FUS and LAF-1 were unable to bind or bundle MTs (**Fig. 2C**). However, fusion of these IDRs to the TPPP-CORE domain to generate to generate synMAP proteins restored MT binding, bundling and condensate formation activities **(Fig. 2D,E and Supplementary Fig. 2c,d).** We quantified the droplet intensity distributions **(Fig. 2F)** and bundling efficiency **(Fig. 2G)** of these synMAPs and compared them to native TPPP-FL and truncations. Across synMAP IDR-CORE chimeras, bundling efficiency (EC50) scaled with droplet formation activity *in vitro* (DDX4>FUS>LAF). The strongest of these synMAPs, DDX4-CORE, had a MT-bundling activity (EC_50_ = 1.47 μM) within 3-fold of wildtype and outperformed many native bundling-competent TPPP-truncations. In contrast, synMAPs based on LAF-1 and the mutant FUSY27S^37^, which only weakly form droplets, showed weak MT-bundling activity **(Fig. 2F,G)**. Thus, synthetic modulation of TPPP condensate formation can control and tune its MT bundling activity analogous to its native IDRs.

While these assays show that synMAP TPPP-CORE-IDR fusions can stimulate microtubule bundling, they cannot resolve ultrastructural differences between the resulting bundled architectures. We thus examined the structure of native and synthetic TPPP microtubule bundles by negative stain electron microscopy. As has been reported previously, TPPP-FL bundles microtubules into long arrays of tightly aligned microtubules^27^. In contrast, no bundling was observed for the condensate-deficient TPPP-CORE **(Fig. 2B)**. synMAP IDR-CORE chimeras produced microtubule bundles in EM that agreed with their quantitative behavior in our *in vitro* bundling assays. DDX4-CORE (EC_50_∼ 1.47 μM) produced bundles that closely resembled those seen for TPPP-FL, creating densely packed structures in which multiple microtubules were aligned together **(Fig. 2D)**. The weaker (EC_50_∼ 5.92 μM) FUS-CORE produced more disperse and less tightly-packed microtubule bundles^38^ **(Fig. 2D)**. No bundled structures were seen for LAF-1-CORE, although some protein clusters were detected on the surfaces of single microtubules **(Fig. 2D)**.

Our results show that TPPP bundling activity can be synthetically controlled using alternative condensate-forming IDR sequences *in vitro* to construct synMAP proteins. The robust correlation we observed between condensate formation and MT bundling across native and synthetic TPPP variants suggests that regulating condensate formation could provide a general mechanism for controlling MT bundling activity in cellular contexts.

### Synthetic clustering of TPPP triggers MT binding, but not bundling, in living cells

Our *in vitro* results suggest that TPPP activity in living cells might be synthetically regulated by controlling when and where it is allowed to condense. To explore this systematically, we built a blue-light triggered TPPP clustering system in 3T3 cells to reconstitute TPPP activity from the bottom-up **(Fig. 3A)**. Our approach is based on the established “OptoDroplets” system for triggering condensation of IDRs in living cells via Cry2 clustering^39,40^. We first validated this tool by expressing either mCh-Cry2WT or FUS-mCh-Cry2WT as a negative and positive control respectively **(Supplementary Fig. 3a)**. As expected, mCh-Cry2WT showed no clusters upon light stimulation, while FUS-mCh-Cry2WT formed spherical drops distributed throughout the cell^40^. Importantly, these controls showed no cross-reactivity of OptoDroplets with the microtubule cytoskeleton as visualized by the live-cell stain SiR-tubulin.

**Fig. 3.**
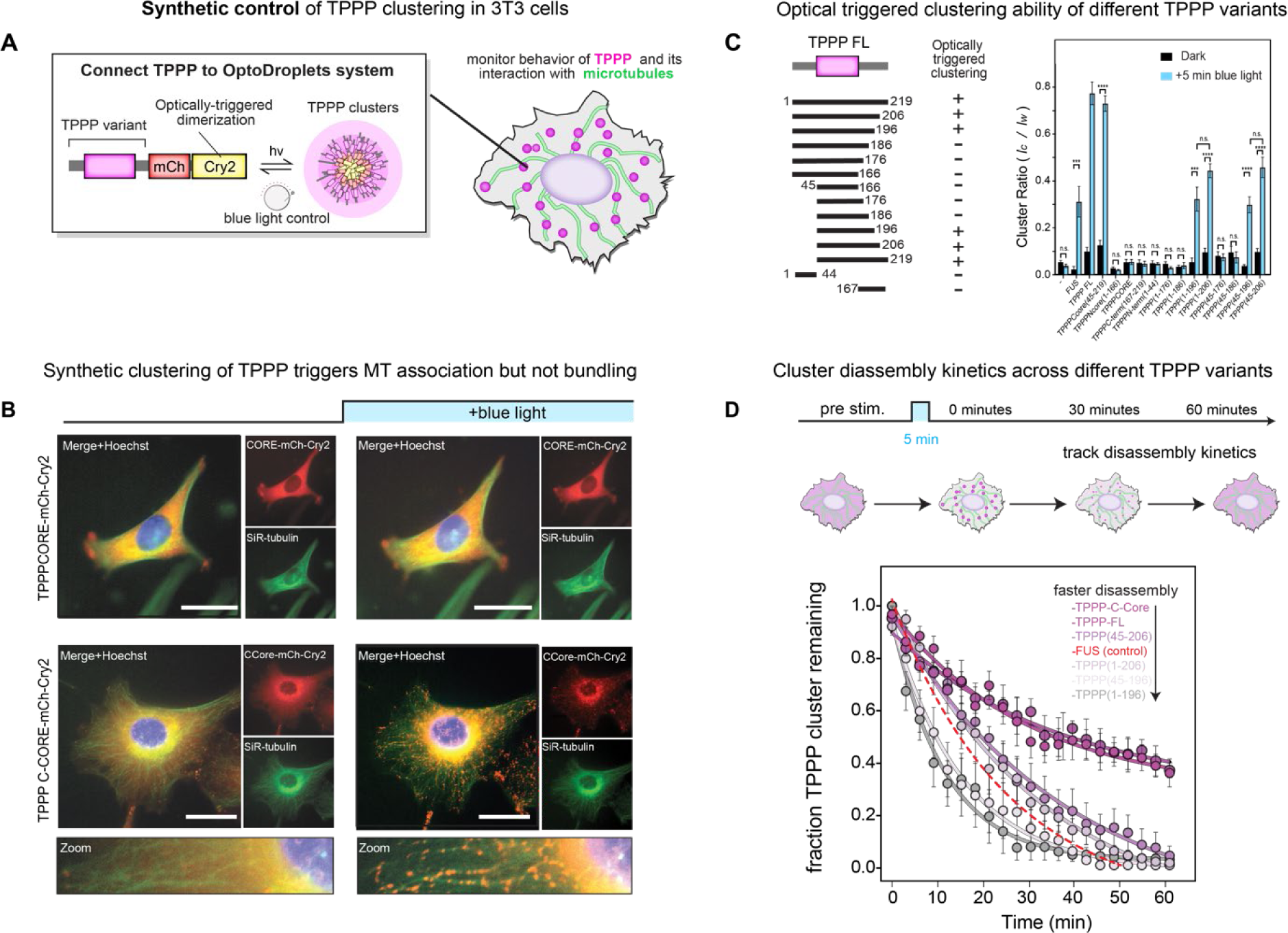
Synthetic clustering of native TPPP fragments triggers microtubule binding, but not bundling, in cells. (A) Schematic diagram of constructs for blue-light triggered clustering of a TPPP fragment in cells by fusion to Cry2WT-mCh. (B) Representative images pre- and post-5 min blue light activation for optical clustering of TPPP-CORE and the C-terminally IDR containing variant C-CORE in NIH3T3 cells (SiR-tubulin, green; Hoechst, blue). Scale bars, 20 μm. (C) Schematic overview and quantification of light-induced cluster formation for different TPPP variants in cells. Clustering activity was scored by dividing cluster intensity (*I_C_*) by the total fluorescence (*I_W_*) of the whole cell. Values are expressed as means ± SEM. Cry2WT only, n = 12; FUS, n = 13; TPPP FL, n = 15; TPPPCcore(45-219), n = 15; TPPPNcore(1-166), n = 15; TPPP(1-206), n = 15; TPPP(45-206), n = 15; TPPP(1-196), n = 12; TPPP(45-196), n = 17; TPPP(1-186), n = 12; TPPP(45-186), n = 14; TPPP(1-176), n = 17; TPPP(45-176), n = 13; TPPPCORE(45-166), n = 12; TPPPN-term(1-44), n = 12; TPPPC-term(167-219), n = 12. Two-tailed Student t test; statistical differences: ****, P < 0.0001; *** P < 0.001; n.s., not significant. (D) Cluster disassembly kinetics in cells across different TPPP variants. Clusters were formed using a five-minute pulse of blue light. Following stimulation, the fraction of clusters remaining over time was tracked by normalization to this initial post-stimulation value. Values are expressed as means ± SEM. FUS, n = 6; TPPP FL, n = 6; TPPPCcore(45-219), n = 7; TPPP(1-206), n = 7 ; TPPP(45-206), n = 7; TPPP(1-196), n = 6; TPPP(45-196), n = 6. The curves were determined by fitting to a one-phase decay equation.

We then applied this tool to study the behavior of different TPPP fragments upon light-induced clustering. Prior to blue-light stimulation, droplet-competent TPPP-FL showed a diffuse distribution within the cell with only faint microtubule decoration. However, upon 5 minutes of blue-light stimulation, TPPP FL formed clusters that associated with and decorated the MTs lattice of the cell **(Supplementary Fig. 3b,c and Supplementary Video S2)**. In contrast, the condensate-deficient TPPPCORE-mCh-Cry2WT remained diffuse upon blue-light stimulation and no cluster formation was observed **(Fig. 3B and Supplementary Video S3)**. TPPP fragments can thus be synthetically clustered to trigger association with MTs, but this does not appear to give rise to extensive bundling in a cellular context^40^.

Applying this opto-clustering method to the panel of TPPP fragments we tested *in vitro*, we found that C-terminal TPPP fragments showed robust cluster-induced MT association in living cells. In contrast, N-terminal TPPP fragments showed no light-dependent clustering onto MTs **(Fig. 3B and Supplementary Fig. 3b)**. Although these N-terminal fragments showed bundling and binding activity *in vitro*, their activity was weaker than C-terminal fragments. This suggests that the requirements for TPPP droplet-regulated interaction with MTs in a living cell may be more stringent than in a minimal *in vitro* system. From this analysis, we defined a minimal fragment (TPPP 45-196) that retains cluster-dependent MT association for synthetic applications **(Fig. 3C).**

This experimental setup also allowed us to investigate how the lifetime of TPPP clusters could be tuned by incorporating different TPPP fragments into our designs. We stimulated cells with blue light for 5 min to assemble TPPP clusters, and then monitored cluster disassembly over time in in the dark **(Fig. 3D and Supplementary Fig. 4)**. Consistent with previous reports, optically induced FUS-mCherry-Cry2 control droplets completely disassembled within 30 minutes in the dark^39^. In contrast, optically clustered TPPP-FL showed slow disassembly, with almost half of all TPPP clusters remaining after one hour **(Fig. 3D and Supplementary Fig. 4)**. Applying this technique to our panel of TPPP variants, we found TPPP disassembly kinetics scaled inversely with *in vitro* and in cell droplet-formation activity. Collectively, these observations suggest that the condensate-forming activity of TPPP we observe *in vitro* can tune its cluster-induced association with microtubules in cells, but that other mechanisms are required to induce formation of higher-order structures.

### synMAP TPPP-IDR condensates generate robust and tunable synthetic microtubule architectures in living cells

Our OptoDroplets experiments revealed that light-induced clustering of TPPP allows it to synthetically associate with MTs in cells. However, this approach did not produce the large-scale MT bundles observed *in vitro* and in the oligodendrocyte cytoskeleton. We noticed that the number of TPPP clusters generated using the OptoDroplets system was high and potentially too finely dispersed to coalesce into higher-order architectures on the timescales accessible to light-stimulation. By comparison, the proposed mechanism for controlling TPPP activity in oligodendrocytes depends on its targeting to a recently discovered class of discrete and highly localized cell-type specific structures called Golgi outposts^27,41–43^. These outposts are thought to act as hubs that concentrate TPPP with interaction partners, including the condensate-forming Golgi matrix protein GM130^27,44,45^.

Inspired by this model, we hypothesized that we could synthetically drive TPPP MT bundling in cells by directly connecting it to synthetic droplet-forming hubs. To develop these hubs, we took advantage of previous work showing that PixD/PixE proteins from *Synechocystis* can form persistent, micrometer-sized liquid-like droplets when fused with IDR sequences^46–48^. We hypothesized that fusion of TPPP and IDR-PixD/E sequences would generate a synMAP-IDR composite of microtubule-binding domains (MTBDs) and condensate-forming sequences (IDRs) that might facilitate microtubule bundling **(Fig. 4A)**.

**Fig. 4.**
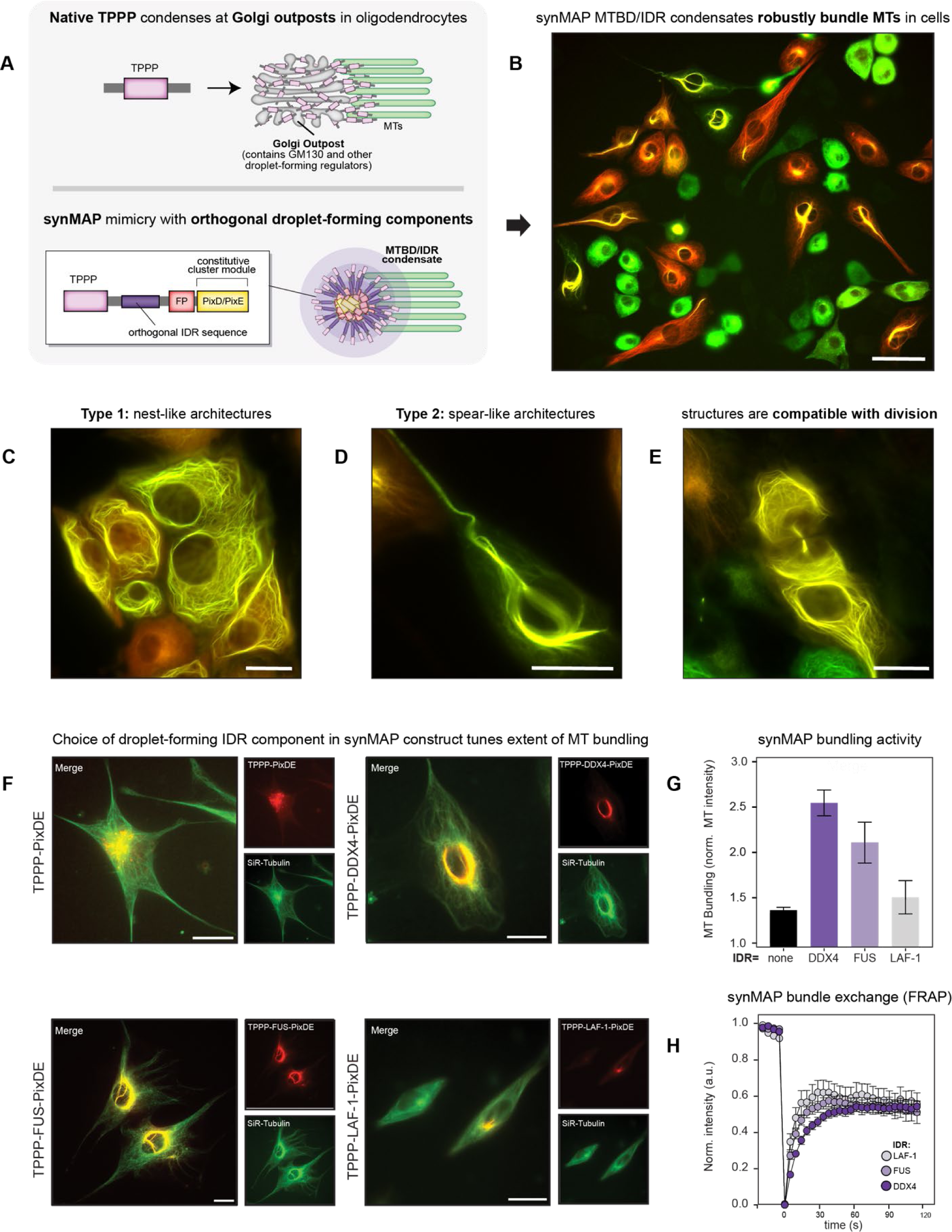
synMAP TPPP-IDR condensates generate robust and tunable synthetic microtubule architectures in living cells. (A) Schematic diagram for the design of synMAP TPPP-IDRs-FP-PixDE constructs. These are engineered to mimic the discrete and localized targeting of native TPPP to Golgi outposts in oligodendrocytes. (B) Representative image of a population of 3T3 cells expressing TPPP-FUS-FusionRed-PixD and TPPP-FUS-Citrine-PixE. SynMAPs condensates organize a diverse range of synthetic microtubule bundling structures across various expression levels. Scale bar, 60 μm. (C) Example image of a TPPP-FUS-PixDE synMAP driven nest-like MT architecture. Images are shown as a merge of the TPPP-FUS-FusionRed-PixD and TPPP-FUS-Citrine-PixE channels. Scale bars, 20 μm. (D) Example image of a TPPP-FUS-PixDE synMAP driven spear-like architecture. Images are shown as a merge of the TPPP-FUS-FusionRed-PixD and TPPP-FUS-Citrine-PixE channels. Scale bars, 20 μm. (E) Example image showing a recently divided cell where a bundled MT-architecture has reformed at the site of division, indicating synMAP structures are compatible with mitosis. Images are shown as a merge of the TPPP-FUS-FusionRed-PixD and TPPP-FUS-Citrine-PixE channels. Scale bars, 20 μm. (F) Example epifluorescence images of microtubule architectures driven by synMAP TPPP-IDR condensates that differ in the identity of the condensate-forming IDR sequence. TPPP-PixDE (negative control), TPPP-DDX4-PixDE, TPPP-FUS-PixDE and TPPP-LAF-1-PixDE images are shown as merged images of the synMAP (red) and SiR-tubulin (green) channels. Scale bars, 20 µm. (G) Quantification of synMAP bundling activity generated by the constructs in (F), defined by the normalized intensity of SiR-tubulin signal near the cell center. Data for TPPP-PixDE (control) or different synMAP TPPP-IDR condensates are shown (mean ± SEM, TPPP-PixDE, n = 3; TPPP-DDX4-PixDE, n = 3; TPPP-FUS-PixD, n = 3; TPPP-LAF-1-PixDE, n = 3). (H) Quantification of exchange dynamics within different synMAP-driven MT bundles, derived from FRAP recovery curves acquired by confocal microscopy. Points within the curve reflect the mean ± SEM: TPPP-DDX4-PixDE, n = 13; TPPP-FUS-PixDE, n = 14; TPPP-LAF-1-PixDE, n = 14. The dynamic recovery curves are derived from non-linear curve fitting based on the one-phase association equation model. Related to (Supplementary Figure 5.)

Expression of TPPP alone did not lead to microtubule bundling in 3T3 cells, and expression of IDR-PixD/E alone produced constitutive droplets dispersed all throughout the cell with no apparent MT association. Remarkably, expression of TPPP-IDR-PixD/E synMAP fusion proteins led to robust and constitutive assembly of large synthetic microtubule architectures in living cells **(Fig. 4B)**. For the TPPP-FUSN-PixE/D synMAP, these composites adopted several visually striking geometries within the cell. These included “nest” like structures encircling the nucleus that spread to the cell cortex; and “spear” like structures that ran from the center of the cell outwards, threading in to extensions of the cell’s overall geometry **(Fig. 4C,D)**. When confined in these narrow spaces, these synthetic architectures would often buckle and twist to adopt more helical shapes. Surprisingly, these synthetic microtubule architectures were fully compatible with cell growth and division, as they were disassembled at the onset of mitosis and rapidly reassembled post cell division **(Fig. 4E and Supplementary Video S5)**. This enabled us to easily generate stable cell lines harboring these novel MT architectures that could be readily maintained across multiple passages.

Our design strategy enabled us to test the ability of other condensate-forming IDR sequences to support synMAP bundle formation in cells. We found that an IDR sequence was absolutely required for bundle formation, as TPPP-PixD/E fusions did not generate bundles. In contrast, DDX4, FUSN, and LAF-1 all produced microtubule bundles in cells, albeit to differing degrees **(Fig. 4F)**. By quantifying the magnitude of MT bundling across multiple cells within the population (detailed in methods), we found that IDRs with stronger droplet-forming activity led to more potent synMAP bundling activity **(Fig. 4G)**. The resulting bundles also showed different exchange dynamics, as judged by their FRAP recovery kinetics, with weaker IDR sequences showing faster exchange **(Fig. 4H and Supplementary Fig. 5)**. Taken together, our results indicate that engineered synMAPs that fuse TPPP to constitutive droplet-forming activity provide a simple and tunable means of robustly generating artificial biologically-compatible MT bundles in mammalian cells.

### Engineered circuits can control synthetic microtubule architectures by regulating the connection between a synMAP’s TPPP and IDR components

Our above results show that synMAPs that directly fuse TPPP to droplet-forming sequences drive constitutive assembly of higher-order microtubule architectures in cells. Thus, we reasoned that synMAPs could be used to build more complex cytoskeletal circuits by separating the TPPP and IDR components and placing their physical connection with one another under regulatable control. To test this idea, we fused one half of the rapamycin-dependent dimerization system FRB to the IDR-PixDE component and fused the other half FKBP to the TPPP component^49,50^. This synMAP circuit design should enable TPPP to be targeted to pre-formed droplet hubs upon rapamycin addition and trigger formation and assembly of synthetic microtubule architectures **(Fig. 5A)**.

**Fig. 5.**
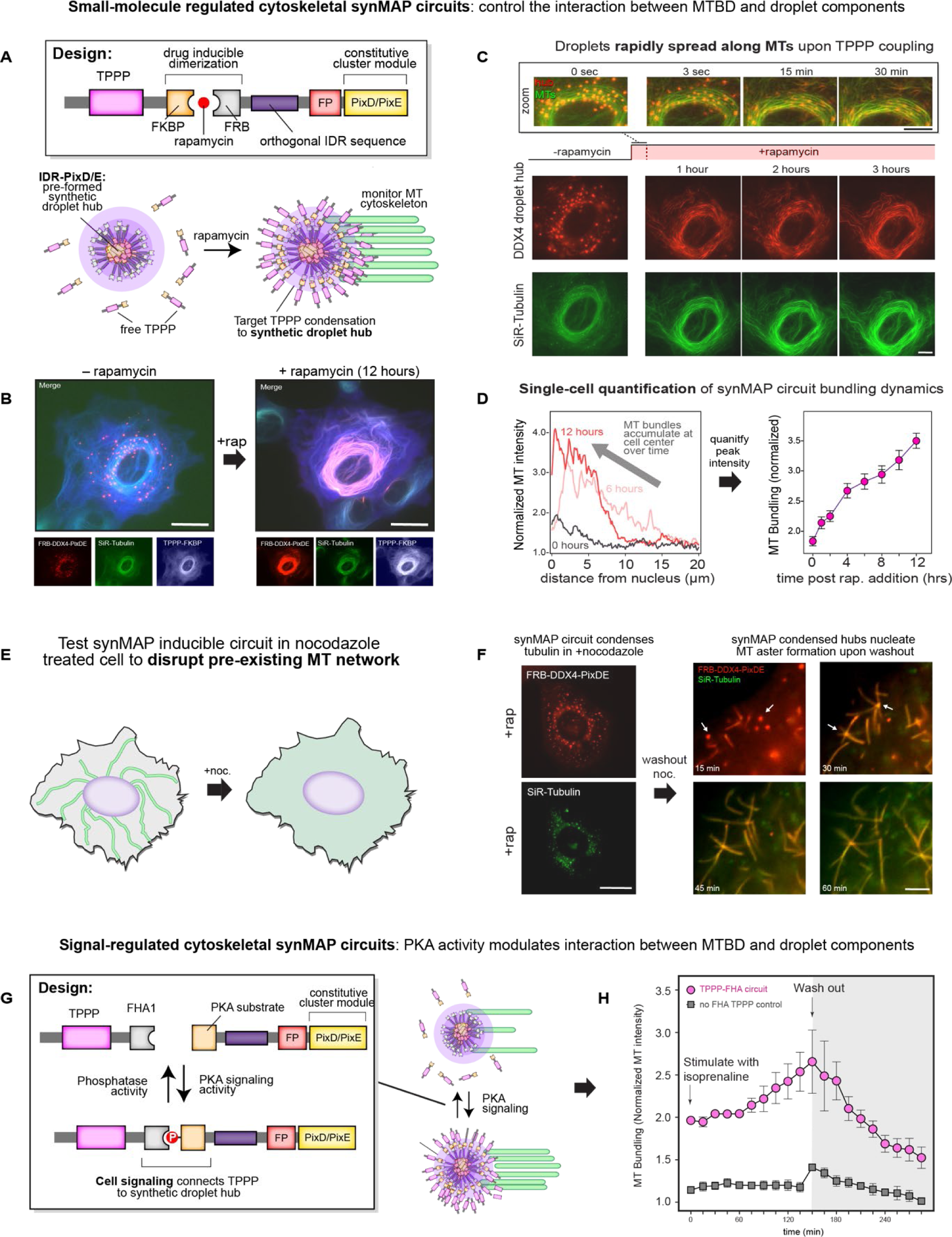
Engineered circuits can control synthetic microtubule architectures by regulating the connection between a synMAP’s TPPP and IDR components. (A) Schematic of the constructs used for building circuits that inducibly connect TPPP to IDR-droplet hubs, mimicking the discrete and localized targeting of native TPPP to Golgi outposts in oligodendrocytes. The system is based on the rapamycin-inducible dimerization pair FKBP/FRB. FRB-IDRs are fused to PixD/E containing fluorescent proteins (FusionRed and Citrine) to produce constitutive droplets, and FKBP is fused to TPPP tagged with tagBFP. Upon rapamycin treatment, a synMAP’s MT binding and IDR components become connected to stimulate assembly of higher-order MT structures. (B) Representative images of NIH3T3 cells co-transduced with an inducible synMAP circuit (FRB-DDX4-PixDE and TPPP-FKBP) before and after (12 hr) rapamycin treatment. The addition of rapamycin (20 µM) rapidly translocated the TPPP component (tagBFP, white color in image) into DDX4 condensates (FusionRed, red) to initiate microtubule bundling (SiR-tubulin, green). Scale bar, 20 µm. (C) Images from a time series of the rapamycin-induced synMAP circuit from (B). Microtubule lattices were directly visualized with SiR-tubulin (green). Zoom view (upper) shows the time series for early stages (0-30 min) following rapamycin induction, in which connection of TPPP fragments to condensates causes deformation and wetting of synMAPs onto microtubules Over several hours, these structures coalesce into larger bundled architectures (bottom). Scale bar, 10 µm. (see also Supplementary Video S4). (D) Quantification of the rapamycin-inducible synMAP circuit in (B)-(C) over time. (left) Line profiles showing MT distribution (normalized SiR-tubulin fluorescence) as a function of the distance from the cell center show how microtubules are organized into bundled architectures near the cell-center over time. The line profile for each time point is used to generate a trajectory (right) showing the single-cell MT bundling dynamics (normalized intensity of SiR-tubulin signal at the cell center) over time. Data are shown as (mean ± SEM, n = 24). (E) Schematic for studying synMAP circuit dynamics following nocodazole-induced microtubule depletion. (F) Inducible synMAP circuits (TPPP-DDX4 design) can nucleate microtubule aster formation following nocodazole treatment as shown in (E). (left) Representative image showing SiR-tubulin colocalization with synMAP (FusionRed, red) droplets in cells treated with nocodazole (30 µM) for six hours. No microtubules are present at this time. Scale bar, 20 µm. (right) Live-cell images of synMAP circuit behavior following nocodazole washout. Images show microtubule aster formation (arrows) originating from synMAP TPPP-DDX4 condensates over time. Scale bar, 5 µm. (G) Schematic for a synMAP circuit in which microtubule architecture is regulated by Protein Kinase A (PKA) signaling activity. A PKA substrate was fused to the condensate-forming DDX4-PixDE component and co-expressed with a TPPP-FHA1 (binds phosphorylated PKA substrate) or TPPP (negative control) component in NIH3T3 cells. When the PKA substrate is phosphorylated in response to cell-signaling inputs, TPPP will be connected to the IDR component to stimulate activity. (H) Demonstration of the PKA-regulated synMAP circuit design from (G). Cells were stimulated with isoprenaline and imaged for 135 minutes. Isoprenaline was then washed out and the cells were further imaged for another 135 minutes. The circuit’s bundling dynamics were quantified as in (D). Data are shown as (mean ± SEM, PKAsub-DDX4-PixDE/TPPP-FHA1, n = 3; PKAsub-DDX4-PixDE/TPPP, n = 3)

We first examined the performance of a synMAP circuit targeting TPPP-FL-FKBP to FRB-DDX4-PixDE droplet hubs in 3T3 cells. Prior to addition of rapamycin, TPPP FL-FKBP was distributed throughout the cytoplasm and weakly decorated MTs at the nuclear periphery^51^. Strikingly, rapamycin-induced targeting of TPPP-FL-FKBP to FRB-DDX4-PixDE droplet hubs drove assembly and extensive MT bundling within the cell **(Fig. 5B)**. Detailed inspection of this process revealed that droplet hubs were the initial sites of microtubule interaction **(Fig. 5C)**. Droplets began to deform within seconds of TPPP coupling and completely spread across the microtubule surface within 15 minutes. Over twelve hours, interactions between adjacent TPPP-coated microtubules resulted in extensive redistribution of the cell’s MT network into nest-like bundles in which the SiR-tubulin signal increased >3 fold relative to the start of the experiment **(Fig. 4C,D and Supplementary Video S4)**. Thus, inducible synMAP circuits targeting TPPP to a DDX4-droplet hub allowed temporal control over the formation of synthetic bundle architectures.

TPPP has also been proposed to play a role in the nucleation of microtubules at Golgi outpost hubs in oligodendrocytes^27,42,52^. However, in 3T3 cells, microtubule nucleation through the centriole dominates the cell’s interphase MT network. Thus, to test whether synMAP circuits that target TPPP to a droplet-hub could stimulate aster formation, we first treated cells with nocodazole to depolymerize all microtubules **(Fig. 4E)**. We then targeted TPPP to PixDE-DDX4 droplets using rapamycin and performed a wash-out experiment^52^. Prior to wash-out, TPPP and tubulin strongly colocalized with DDX4 droplet hubs, but no microtubules were detected owing to the presence of nocodazole **(Fig. 4E,F)**. However, upon nocodazole wash-out, MT filaments rapidly emerged from many TPPP droplet-hubs in an aster-like structure **(Fig. 4F and Supplementary Video S6)**. Over time, multiple droplet-asters clustered and fused together to form MT bundles that coalesced into nest-like architectures as described earlier **(Supplementary Fig. 6 and Supplementary Video S6)**. This result indicates that synMAP circuits can stimulate and stabilize the nucleation of new MTs when sufficiently isolated from other sources of nucleation and polymerization.

Next, we tested whether the same design principles governing the FKBP/FRB synMAP circuit could be used to modulate microtubule architectures in response to endogenous cell-signaling activity, using Protein Kinase A (PKA) signaling as a test case. To this end, we replaced the FKBP/FRB heterodimerization module with an FHA/PKAsub interaction pair, in which the FHA domain only binds to PKAsub when it is phosphorylated by PKA^53,54^. Expression of this TPPP-FHA1 / PKAsub-DDX4-PixD/E synMAP circuit led to moderate levels of constitutive bundling in unstimulated cells, consistent with some basal interaction between the FHA1 and PKAsub components^55^ **(Fig. 5G)**. Stimulation of these cells with a saturating dose of the PKA agonist isoprenaline led to a quantitative increase MT bundling over two hours, and decreased back to baseline levels within one hour following wash out **(Fig. 5G and Supplementary Video S7).** In contrast, a TPPP / PKAsub-DDX4-PixD/E negative control circuit that lacks the FHA1 interaction module showed neither baseline nor inducible MT bundling **(Fig. 5G, Supplementary Fig. 7).**

Together, these synMAP circuits demonstrate that the formation and dynamics of a synMAP microtubule architecture can be controlled by regulating the connection between the microtubule binding components and the droplet-forming IDR components. Thus, synMAPs can be effectively used as a module for more complex circuit designs in mammalian cells.

### Microtubule interaction strength and IDR potency provide a two-level control scheme for designing inducible synMAP bundling circuits in cells

In the above synMAP circuits, we found that inducibly targeting TPPP to a DDX4 droplet hub led to extensive bundling activity, which could be quantified by tracking accumulation of MTs towards the perinuclear region of the cell over time. These metrics allow us to take a systematic approach to experimentally define how bundling activity of a synMAP circuit is quantitatively affected by either 1) the strength of the TPPP microtubule binding component or 2) the strength of the droplet-forming IDR component **(Fig. 6A,E)**.

**Fig. 6.**
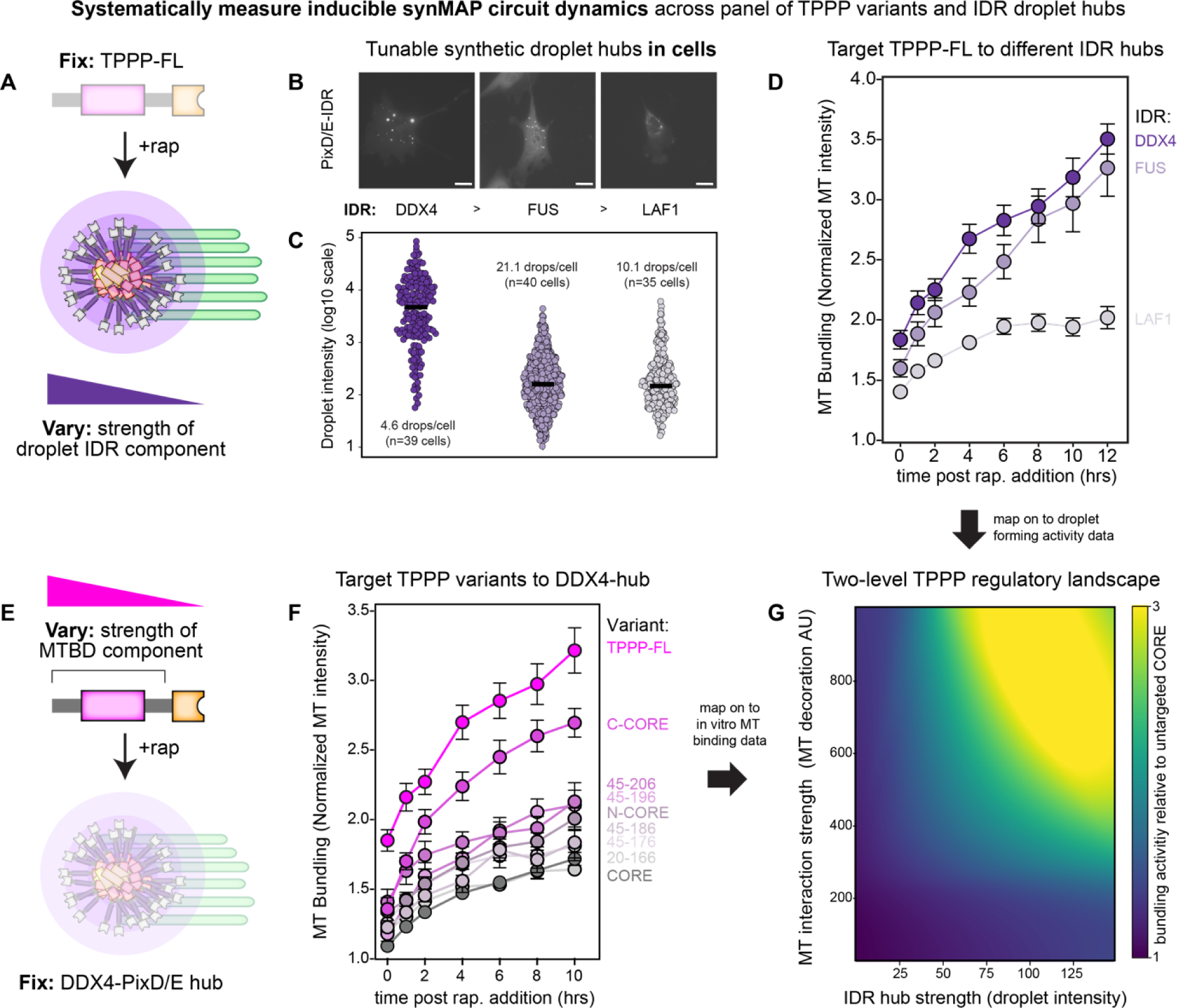
Microtubule interaction strength and IDR potency provide a two-level control scheme for tuning inducible synMAP bundling circuits in cells. (B) Schematic depicting strategy for systematically varying the condensate-forming component (IDR sequence) of an inducible synMAP TPPP-IDR circuit. (C) Representative images from NIH3T3 cells (FusionRed, white color in images) showing the size and distribution of the IDR-droplet component of the different synMAP circuits. Scale bar, 20 µm. (D) Quantification of the IDR-droplet components from (B), based on FusionRed fluorescence signal intensity in NIH3T3 cells. Median intensity is indicated from measurements from 39, 40, and 35 representative cells, with total droplet numbers respectively: FRB-DDX4-PixDE, n = 182; FRB-FUS-PixDE, n = 843; FRB-LAF-1-PixDE, n = 354. From these data, an average number of droplets per cell was also determined as: FRB-DDX4-PixD = 4.1 drops/cell, FRB-FUS-PixDE = 21.1 drops/cell, and FRB-LAF-1-PixDE = 10.1 drops/cell. (E) Quantification of synMAP circuit induction for the panel of IDR-varying designs from (A), using the method described in Fig. 5D. Data are shown as (mean ± SEM, FRB-DDX4-PixDE/TPPP-FL-FKBP, n = 24; FRB-FUS-PixDE/TPPP-FL-FKBP, n = 27; FRB-LAF-1-PixDE/TPPP-FL-FKBP, n = 31). Related to (Supplementary Figure 8.) (F) Schematic depicting strategy for systematically varying the microtubule-interaction component (TPPP variant) of an inducible synMAP TPPP-IDR circuit. (G) Quantification of synMAP circuit induction for the panel of TPPP-varying designs from (E), using the method described in Fig. 5D. Data are shown as (mean ± SEM, TPPP-FL-FKBP/FRB-DDX4-PixDE, n = 24; TPPPCcore-FKBP/FRB-DDX4-PixDE, n = 16; TPPPNcore-FKBP/FRB-DDX4-PixDE, n = 15; TPPP(45-206)-FKBP/FRB-DDX4-PixDE, n = 12; TPPP(45-196)-FKBP/FRB-DDX4-PixDE, n = 12; TPPP(45-186)-FKBP/FRB-DDX4-PixDE, n = 18; TPPP(45-176)-FKBP/FRB-DDX4-PixDE, n = 26; TPPP(20-166)-FKBP/FRB-DDX4-PixDE, n = 15; TPPP-CORE-FKBP/FRB-DDX4-PixDE, n = 36). Related to (Supplementary Figure 9.) (H) Two-level regulatory landscape for inducible synMAP circuit dynamics. Dynamic bundling metrics from (D) and (F) were mapped onto a space defined by a TPPP variant’s microtubule binding affinity (from single-MT binding assay) and the strength of the IDR hub (defined by large-droplet intensity). The data were interpolated to create a phenomenological phase portrait describing the inducible synMAP circuit’s dynamics as a function of the two parameters.

To compare the effect of different IDR droplet-hubs, we first expressed different FRB-IDR-PixDE constructs in 3T3 cells and characterized the size, number, and exchange dynamics of the resulting hubs. We found that FRB-DDX4-PixDE created the greatest number of large droplets (>1000 a.u.) in cells, followed by FRB-FUS-PixDE (large number of small droplets) and FRB-LAF-1-PixDE (small number of small droplets) **(Fig. 6B,C)**. We then built synMAP circuits targeting TPPP-FL to each class of IDR droplet-hub and quantified bundle formation at the single-cell level across multiple replicate cells **(Fig. 6A,D)**. When TPPP-FL was targeted to DDX4 droplet-hubs, MT-bundling was strong, producing the highest density of MTs in bundles over a 10-hour period. FUS droplet-hubs also produced substantial MT bundling, showing a similar rate of bundling as DDX4 but with slightly reduced basal activity and a lower end-point. In contrast, the weaker droplet-forming LAF-1 hubs showed significantly lower bundling rates and lower end-point bundling **(Fig. 6D and Supplementary Fig. 8).** These synMAP circuit dynamics correlate with the droplet-formation strength of the IDR-PixD/E module (DDX4>FUS>LAF1) **(Fig. 2)**, indicating that synMAP circuit behavior in cells can be tuned by manipulating the microscale properties of the condensate component.

We next tested the effects of manipulating the strength of the TPPP MT binding component on synMAP circuit behavior **(Fig. 6E)**. Having seen that DDX4 droplet-hubs had the most striking bundling phenotypes, we locked in this component of the circuit and targeted TPPP fragments with different interaction strength (as defined by our *in vitro* and optoDroplets analyses) to these hubs. We found that TPPP variants with strong MT-association showed the strongest microtubule bundling activity when targeted to DDX4-droplet hubs (e.g. C-CORE) **(Fig. 3C**, **Fig. 6F, Supplementary Fig. 2b, and Supplementary Fig 9a-c)**. Variants that could bundle microtubules *in vitro* but showed poor activity in the OptoDroplets assay showed limited activity (e.g. N-CORE) **(Fig. 3C**, **Fig. 6F and Supplementary Fig. 9d-f)**. Targeting TPPP-CORE to DDX4 droplets drove weak, but detectable, MT bundling in this assay **(Supplementary Fig. 9g-i,and Supplementary Video S8)**. However, the degree of rescue is dramatically lower than what we observed *in vitro*. Indeed, the majority of TPPP-CORE remained in droplets when targeted to DDX4 hubs, in contrast to other TPPP variants that induced droplet-spreading along the MT surface. This suggests that although a droplet hub can enhance a weak MT-binding activity, if droplet formation is too strong it may outcompete the target MT interaction in cellular contexts.

Having collected experimental inducible bundling dynamics for synMAP circuits spanning a range of TPPP variants and droplet hubs across more than a hundred different cells, we mapped these metrics onto a phase space defined by a TPPP variant’s MT interaction strength (extent of MT decoration *in vitro*) and strength of the IDR hub (large droplet intensity). Interpolating these data produced a phenomenological design manual for programming synMAP circuit behavior **(Fig. 6G)**. Stronger MT interaction strength allowed for some synMAP-driven bundling even in the absence of direct targeting to a droplet hub. Bundling activity dramatically increased across all variants when targeted to a strong droplet hub (DDX4), but TPPP fragments with reduced MT interaction strength were less effective. Together, this landscape suggests two routes for spatiotemporal control of synMAP circuit activity in cells: modulation at the nanoscale, targeting how strongly TPPP can engage microtubules; or modulation at the microscale, targeting the condensate-formation activity of the IDR component. As a result, different synMAP circuits and behaviors can be designed by choosing different paths through this landscape.

## Discussion

We have identified and exploited a condensate-regulatable microtubule-binding activity from the oligodendrocyte protein TPPP to develop synMAPs: engineered proteins that provide tunable control over higher-order microtubule structure formation *in vitro* and in mammalian cells. The central design feature of synMAPs is the pairing of TPPP’s native microtubule-binding activity with other orthogonal, condensate-forming IDR sequences. *In vitro*, direct fusion of IDRs to TPPP’s minimal structured domain stimulates potent MT bundling that quantitatively correlates with the droplet-forming activity of the IDR. In living cells, synthetic MT bundling requires connection of larger TPPP fragments to constitutive droplet-forming sequences. We capitalized on this stringency to engineer cytoskeletal circuits in cells in which synMAP activity could be controlled by regulating the interaction between the MTBD and droplet-forming components. This allowed the formation and dynamics of microtubule architectures to be synthetically manipulated by small molecules and endogenous cell-signaling activities. By systematically analyzing synMAP circuit behavior using different MTBDs and droplet-forming sequences, we developed a two-level design manual for programming synMAP-triggered assembly of MT architectures at both the nanoscale and microscale.

A surprising result that emerged from our synthetic approach is that many critical cytoskeletal activities—MT binding, bundling, and aster formation—could all be driven and regulated through the action of arbitrary droplet-forming sequences which have not specifically co-evolved with MTs to perform these functions. Indeed, the ability of TPPP’s MT-binding domain to cooperate so effectively with the IDRs of DDX4—an RNA helicase^56,57^—indicates considerable plasticity in the mechanisms by which condensate-formation can synergize with MT organization. We suspect this flexibility may explain the multitude of MT structures other TPPP isoforms are associated with outside of oligodendrocytes, including the sperm flagellum (TPPP2)^25,58^ and protozoan ciliary lattices^59,60^. More generally, our results suggest that droplet-forming sequences may provide a versatile *biophysical* strategy for coordinating the nanoscale interaction between MAPs and MTs across cellular length scales **(Fig. 6G)**.

At the same time, these general-purpose biophysical mechanisms must be paired with more sophisticated regulatory mechanisms to reliably specify and control the assembly of target architectures with specific cellular functions. For example, while TPPP was able to potently bundle MTs within the cell, the interphase MT network derived from centriolar MTOCs dominated the overall structure and spatial distribution of the emergent structure towards the cell center^61–63^. This motivated our embedding of synMAPs in more complex cytoskeletal circuit designs that provided additional temporal control over when bundling could occur. We anticipate that extension of these circuits to incorporate spatial control over synMAP activity could lead to more precise organization and positioning of synthetic MT architectures within the cell. Nevertheless, our results show droplet-based biophysical mechanisms provide a tractable starting point for the design of higher-order microtubule-based cellular machines.

The synMAP circuits we present generate new cytoskeletal structures inside cells, but do not yet encode new biological functions. However, our droplet-based design provides straightforward paths going forward to equip these structures with additional activities based on co-condensation of IDR-matched client proteins into the structure^14,64^. Such strategies could be used to selectively recruit specific motor proteins to generate bending forces and motion, or to localize and position biochemical cargos or protein payloads and activities within the cell. By synthetically building these structures and systems from the ground up, synMAPs will reveal general design principles that provide a deeper ability to understand the diversity of complex microtubule architectures seen in nature and a path forward to engineering new microtubule structures and machines of our own design.

## Acknowledgments

We thank members of the Coyle Lab, A. Weeks, W. Bement, A. Suzuki, and C. Lim for advice, helpful discussions.

## Author contributions

CCC and SMC designed the project; CCC performed all experiments and interpreted the data for the manuscript under the supervision of SMC; CCC and SMC wrote the manuscript and contributed to the final version.

## Methods

### Plasmid construction

Homo sapiens TPPP WT gene was synthesized as gBlocks (Integrated DNA Technologies). TPPP truncation mutants were produced using a standard PCR. All TPPP variants were then inserted into a pBH4 vector which contains GFP sequence and His-tag at N-terminal region. Homo sapiens DDX4 (residues 1-236), LAF-1 RGG (residues 1-200) and FUS Y27S mutant (residues 1-214) fragments were obtained as gBlocks (Integrated DNA Technologies). Human FUS (residues 1-214) was amplified by PCR using pHR-FUSN-mCh-Cry2WT (Addgene #101223). The IDRs fragments were fused with TPPPCORE domain(aa.45-166) into the same pBh4-His-GFP plasmid. For making the OptoTPPP, different TPPP fragment were produced by PCR then cloned into the pHR-mCh-Cry2WT plasmid.

The DNA fragments of FRB, IDRs, FusionRed-PixD and Citrine-PixE were produced by PCR using existing templates (Addgene #31181), (Addgene #111503) and (Addgene #111505) and subcloned into an SFFV vector to generate FRB-IDR-FusionRed-PixD and FRB-IDR-Citrine-PixE. The DNA fragments of TPPP FL and FKBP were produced by PCR using existing templates and (Addgene #31184). The tagBFP gene was synthesized as gBlocks (Integrated DNA Technologies). These fragments were subcloned into an SFFV vector to generate TPPP-tagBFP-FKBP. For making different TPPP variants-tagBFP-FKBP constructs, the DNA fragments were generated by PCR and then fused into the SFFV promoter vector.

To create the TPPP-tagBFP-FHA1 and PkAsub-DDX4-FusionRed-PixD/Citrine-PixE constructs, we first fused the PKA substrate sequence "LRRATLVD" to the N-terminus of DDX4-FusionRed-PixD and DDX4-Citrine-PixE, resulting in the PKAsub-DDX4-FusionedRed-PixD and PKAsub-DDX4-Citrine-PixE constructs. Next, we generated the FHA1 gene via PCR using existing templates (addgene #138202). This FHA1 gene was then introduced to replace the FKBP sequence in the TPPP-tagBFP-FKBP construct, resulting in the creation of the TPPP-tagBFP-FHA1 construct. All fragments were subcloned to the SFFV promoter vector.

### Bacterial protein expression and purification

All TPPP variant and TPPP-chimeras constructs were expressed in BL21(DE3). *Escherichia coli* overnight at 16°C after induction with 0.5 mM IPTG. The culture was harvested by centrifugation and lysed according to the following procedure. The cell pellets were resuspended in lysis buffer (50 mM KH2PO4, 50 mM Na2HPO4, 150 mM NaCl, and 2 mM β-mercaptoethanol), disrupted by French press, and centrifuged at 15,000 g at 4 C for 20 min. The supernatant was incubated with Ni resin (Sigma) for 30 min, followed by a prewash with 25 mM imidazole. Proteins were eluted with the buffer containing 250 mM imidazole. Then the samples were further purified by SEC (Superdex 200 16/60) and analyzed by SDS-PAGE. The peak protein fraction was collected, concentrated, and stored in a buffer consisting of 20 mM Hepes, pH 7.4, 50 mM NaCl, and 3 mM DTT. All proteins were flash frozen and stored at −80 °C. Before use, all proteins were pre-cleared of aggregates via centrifugation at 4 °C.

### Single-microtubule binding and microtubule-bundling assays

Microtubules were polymerized from a solution that contained unlabeled, Alexa Fluor 594-labeled and biotin-labeled tubulin in a ratio of 25:1:1 at 37°C in the presence of GMPCPP. Flow chambers were assembled with biotin-PEG-coated coverslips. The chamber was sequentially filled with 0.5 mg/ml α-casein, 0.2 mg/ml NeutrAvidin, and biotinylated microtubules in 1xBRB80, and 20 μM Taxol. Next, GFP-labeled TPPP variants or TPPP-Chimeras were added to the chamber in 1xBRB80 supplemented with 20 μM Taxol, 0.5 mg/ml α-casein, 10% sucrose, 2 mM DTT, 200 μg/ml glucose oxidase, 35 μg/ml catalase, and 4.5 μg/ml glucose. The chamber was then filled with the solution and sealed, then imaged by using TIRF microscopy. The GFP and Fluor 594 intensities were determined using Fiji.

For a microtubule bundling assay, TPPP variants and TPPP chimeras were incubated with Taxol-stabilized, Alexa Fluor 594-labeled and biotin-labeled tubulin microtubules for 5 minutes. The mixture of bundled and free MTs were captured onto a biotinylated coverslip for quantitative determination of the fraction of MTs bundled. Microtubule bundles were visualized by a TIRF illumination system, and Fluor 594 intensities were determined using Fiji. Single microtubules and bundled microtubules were distinguished by the fluorescence intensity line profile along the structure, using the intensity distribution from an unbundled, single MT sample as a reference. A bundled structure was defined defined as any structure with intensity greater than two standard deviations above the mean-value for the single MT distribution. Using this metric, the bundle-fraction for a given condition was defined as the fraction of MT structures classified as bundles in the sample. This method was used to produce all saturation curves, in which a concentration series was performed for each TPPP variant or TPPP-Chimeras. EC50s were obtained by fitting these curves to a one-site-binding hyperbola^22,31,65^.

### Droplet formation assay

GFP-TPPP variants and TPPP-chimeras with 20 μM in buffer (20 mM HEPES, 50 mM NaCl, 3 mM DTT) containing crowding reagents (12% dextran), respectively. Samples (2 μl) were spotted onto glass slides then visualized using Confocal fluoresce microscopy. Images were visualized GFP signal; then measured and quantified using Fiji.

### Fluorescence recovery after photobleaching (FRAP) analysis

GFP-TPPP droplets were FRAP using confocal microscopy (NIKON A1RS) for a total of 5 min. Defined regions were photobleached and monitored the at the 488 nm wavelength fluorescence intensities; in the NIH3T3 living cell, the TPPP-IDRs-FusionRed/Citrine-PixDE bundled MTs were FRAP and monitored at the 561 nm wavelength fluorescence intensities in every 5 s intervals for a total of 2 min. Recovery fluorescence intensities were recorded for the indicated time then, quantified and normalized by using Fiji.

### Negative stain electron microscopy

The GFP-TPPP FL, GFP-TPPPCORE, and GFP-SynMAPs were mixed with unlabeled GMPCPP-stabilized microtubules in the 1xBRB80 buffer for 5 min at room temperature. The mixed samples were diluted with 1xBRB80 to reduce unbundled microtubules in the background, and a 3 µl mixture was spotted on glow-discharged grids (Electron Microscopy Sciences, CF400-Cu), then stained with 2% uranyl acetate. Images were collected at a magnification of 45,000 using 120 kV Talos L120C at UW-Madison Cryo-EM Research Center.

### Construction of stable cell lines

Optogenetic Cry2WT expressing constructs and FRB-IDRs-PixDE/TPPP variants-FKBP constructs were produced using lentivirus. Pantropic VSV-G pseudotyped lentivirus was produced by transfecting 293T cells (ATCC CRL-3216) with a pLV-EF1a-IRES or pTwist-SFFV transgene expression vector and the viral packaging plasmids psPAX2 and pMD2.G using Fugene HD (Promega #E2312). Viral production was performed in 6-well tissue culture treated plates (Corning 3335). Viral supernatants were collected 3 d after transfection and passed through a 0.45-mm filter to remove cell debris and then incubated with NIH 3T3 cells containing polybrene 8 µg/mL. Viral medium was replaced with normal growth medium 24 hr after infection. After 3 days, cells were sorted on an FACSAria cell sorter (BD Biosciences) based on fluorescent protein expression levels.

### Live cell imaging

Cells were plated on 0.17±5 μm 24 well glass bottom plate (Cellvis) in the FluotoBrite^TM^ DMEM (ThermoFisher A1896701) overnight. All live cell imaging was performed using 60X oil immersion objective (NA 1.4) on a Nikon Ti-Eclipse with a temperature stage at 37°C and supplied with 5% CO2. NIH3T3 cells were imaged using four wavelengths (405 nm for tagBFP; 488 nm for Citrine; 561 nm for mCherry or FusionRed; 674 nm for SiR700-Tubulin Kit, #CY-SC014, Cytoskeleton Inc.).

### OptoDroplet assay

The OptoTPPP was designed based on the strategy from the original OptoDroplets study^39^. NIH3T3 cells were transfected with OptoFUS control constructs and OptoTPPP variants constructs to test. Cells were plated on 0.17±5μm 24 well glass bottom plate (Cellvis) in FluotoBrite^TM^ DMEM (ThermoFisher A1896701) for imaging. Cluster formation was induced using Nikon Ti-Eclipse with a Mightex Polygon digital micromirror device (DMD) at 475 nm wavelength light. The clusters and microtubules were imaged using two wavelengths (561 nm for mCherry; 647 nm for SiR700-tubulin).

To quantify the cluster ratio in cells containing clusters, total cluster intensity (*I_C_*) was divided by the whole cell’s total fluorescence intensity (*I_W_*) adapted from a previous study^66^. The cluster disassembly rate was based on quantifying cluster number following stimulation over time normalized to the initial post-stimulation number. Curves were quantified by fitting them to a one-phase decay equation.

### Quantification of MT bundles in living cells

Microtubule bundles were defined based on the fluorescence intensity of SiR-tubulin using a previously established metohd^67,68^. In each cell, we identified the region with the highest microtubule bundling (often near the cell center of cells) and measured the SiR-tubulin fluorescence intensity along straight-line segments of approximately 5 μm in length. This intensity was then normalized by dividing it by the fluorescence intensity of single microtubules within the same cell, yielding the normalized microtubule bundling intensity^67,68^. All the quantification analyses of the fluorescence images were performed using Fiji.

### Nocodazole treatments

For microtubule depolymerization, nocodazole (30 μM) was added to culture media for 6 hr. Then the NIH3T3 cells were washed with cold FluotoBriteTM medium directly at the microscope with temperature stage at 37°C and supplied with 5% CO2. The heated stage slowly raised the temperature. The regrowth microtubule images and aster formation were captured using Nikon Ti-Eclipse at the wavelengths indicated in the figures.

### Statistical analysis

Statistical analyses were calculated using Prism (version 9.0c). Figure legends detail the n values and error bars for each experiment. Statistical differences for single-MT binding assay (Supplementary Figure 2c,e) and cluster formation assay (Fig. 3C) were determined by Student’s two-tailed t-test; *P< 0.05, **P< 0.01, ***P< 0.001 and ****P< 0.0001.

## References

1. Vale, R. D. The Molecular Motor Toolbox for Intracellular Transport. Cell 112, 467–480 (2003).

2. Nogales, E. Structural insights into microtubule function. Annu. Rev. Biophys. Biomol. Struct. 30, 397–420 (2001).

3. Olmsted, J. B. Microtubule-Associated Proteins. Annu. Rev. Cell Biol. 2, 421–457 (1986).

4. Haimo, L. T. & Rosenbaum, J. L. Cilia, flagella, and microtubules. J. Cell Biol. 91, 125s–130s (1981).

5. Goodson, H. V. & Jonasson, E. M. Microtubules and Microtubule-Associated Proteins. Cold Spring Harb. Perspect. Biol. 10, a022608 (2018).

6. Bodakuntla, S., Jijumon, A. S., Villablanca, C., Gonzalez-Billault, C. & Janke, C. Microtubule-Associated Proteins: Structuring the Cytoskeleton. Trends Cell Biol. 29, 804–819 (2019).

7. Roybal, K. T. & Lim, W. A. Synthetic Immunology: Hacking Immune Cells to Expand Their Therapeutic Capabilities. Annu. Rev. Immunol. 35, 229–253 (2017).

8. Morsut, L. et al. Engineering Customized Cell Sensing and Response Behaviors Using Synthetic Notch Receptors. Cell 164, 780–791 (2016).

9. Roybal, K. T. et al. Engineering T Cells with Customized Therapeutic Response Programs Using Synthetic Notch Receptors. Cell 167, 419–432.e16 (2016).

10. Markson, J. S. & Elowitz, M. B. Synthetic Biology of Multicellular Systems: New Platforms and Applications for Animal Cells and Organisms. ACS Synth. Biol. 3, 875–876 (2014).

11. Chaaban, S. & Brouhard, G. J. A microtubule bestiary: structural diversity in tubulin polymers. Mol. Biol. Cell 28, 2924–2931 (2017).

12. LaFrance, B. J., et al. Structural transitions in the GTP cap visualized by cryo-electron microscopy of catalytically inactive microtubules. Proc. Natl. Acad. Sci. 119, e2114994119 (2022).

13. Saeidi, H. R., Lohrasebi, A. & Mahnam, K. External electric field effects on the mechanical properties of the αβ-tubulin dimer of microtubules: a molecular dynamics study. J. Mol. Model. 20, 2395 (2014).

14. Wang, B. et al. Liquid–liquid phase separation in human health and diseases. Signal Transduct. Target Ther. 6, 290 (2021).

15. Hyman, A. A., Weber, C. A. & Jülicher, F. Liquid-Liquid Phase Separation in Biology. Annu Rev Cell Dev Bi 30, 39–58 (2014).

16. Li, P. et al. Phase transitions in the assembly of multivalent signalling proteins. Nature 483, 336– 340 (2012).

17. Brangwynne, C. P. et al. Germline P Granules Are Liquid Droplets That Localize by Controlled Dissolution/Condensation. Science 324, 1729–1732 (2009).

18. Hernández-Vega, A. et al. Local Nucleation of Microtubule Bundles through Tubulin Concentration into a Condensed Tau Phase. Cell Rep. 20, 2304–2312 (2017).

19. King, M. R. & Petry, S. Phase separation of TPX2 enhances and spatially coordinates microtubule nucleation. Nat. Commun. 11, 270 (2020).

20. Sun, M. et al. NuMA regulates mitotic spindle assembly, structural dynamics and function via phase separation. Nat. Commun. 12, 7157 (2021).

21. Song, X. et al. Phase separation of EB1 guides microtubule plus-end dynamics. Nat. Cell Biol. 25, 79–91 (2023).

22. Chang, C.-C., Huang, T.-L., Shimamoto, Y., Tsai, S.-Y. & Hsia, K.-C. Regulation of mitotic spindle assembly factor NuMA by Importin-β. J. Cell Biol. 216, 3453–3462 (2017).

23. Chang, C.-C. & Hsia, K.-C. More than a zip code: global modulation of cellular function by nuclear localization signals. FEBS J. 288, 5569–5585 (2021).

24. Alfaro-Aco, R., Thawani, A. & Petry, S. Structural analysis of the role of TPX2 in branching microtubule nucleation. J. Cell Biol. 216, 983–997 (2017).

25. Vincze, O. et al. Tubulin Polymerization Promoting Proteins (TPPPs): Members of a New Family with Distinct Structures and Functions. Biochemistry 45, 13818–13826 (2006).

26. DeBonis, S., Neumann, E. & Skoufias, D. A. Self protein-protein interactions are involved in TPPP/p25 mediated microtubule bundling. Sci. Rep. 5, 13242 (2015).

27. Fu, M. et al. The Golgi Outpost Protein TPPP Nucleates Microtubules and Is Critical for Myelination. Cell 179, 132–146.e14 (2019).

28. Oláh, J. & Ovádi, J. Pharmacological targeting of α-synuclein and TPPP/p25 in Parkinson’s disease: challenges and opportunities in a Nutshell. FEBS Lett. 593, 1641–1653 (2019).

29. Oláh, J., Bertrand, P. & Ovádi, J. Role of the microtubule-associated TPPP/p25 in Parkinson’s and related diseases and its therapeutic potential. Expert Rev. Proteomic 14, 301–309 (2017).

30. Ubhi, K., Low, P. & Masliah, E. Multiple system atrophy: a clinical and neuropathological perspective. Trends Neurosci. 34, 581–590 (2011).

31. Hsia, K.-C. et al. Reconstitution of the augmin complex provides insights into its architecture and function. Nat. Cell Biol. 16, 852–863 (2014).

32. Patel, A. et al. A Liquid-to-Solid Phase Transition of the ALS Protein FUS Accelerated by Disease Mutation. Cell 162, 1066–1077 (2015).

33. McGurk, L. et al. Poly(ADP-Ribose) Prevents Pathological Phase Separation of TDP-43 by Promoting Liquid Demixing and Stress Granule Localization. Mol. Cell 71, 703–717.e9 (2018).

34. Elbaum-Garfinkle, S. et al. The disordered P granule protein LAF-1 drives phase separation into droplets with tunable viscosity and dynamics. Proc. Natl. Acad. Sci. 112, 7189–7194 (2015).

35. Nott, T. J. et al. Phase Transition of a Disordered Nuage Protein Generates Environmentally Responsive Membraneless Organelles. Mol. Cell 57, 936–947 (2015).

36. André, A. A. M. & Spruijt, E. Liquid–Liquid Phase Separation in Crowded Environments. Int. J. Mol. Sci. 21, 5908 (2020).

37. Lin, Y., Currie, S. L. & Rosen, M. K. Intrinsically disordered sequences enable modulation of protein phase separation through distributed tyrosine motifs. J. Biol. Chem. 292, 19110–19120 (2017).

38. Gouveia, B. et al. Capillary forces generated by biomolecular condensates. Nature 609, 255–264 (2022).

39. Shin, Y. et al. Spatiotemporal Control of Intracellular Phase Transitions Using Light-Activated optoDroplets. Cell 168, 159–171.e14 (2017).

40. Zhang, X. et al. The proline-rich domain promotes Tau liquid–liquid phase separation in cells. J. Cell Biol. 219, e202006054 (2020).

41. Ori-McKenney, K. M., Jan, L. Y. & Jan, Y.-N. Golgi Outposts Shape Dendrite Morphology by Functioning as Sites of Acentrosomal Microtubule Nucleation in Neurons. Neuron 76, 921–930 (2012).

42. Valenzuela, A., Meservey, L., Nguyen, H. & Fu, M. Golgi Outposts Nucleate Microtubules in Cells with Specialized Shapes. Trends Cell Biol. 30, 792–804 (2020).

43. Chung, C. G. et al. Golgi Outpost Synthesis Impaired by Toxic Polyglutamine Proteins Contributes to Dendritic Pathology in Neurons. Cell Rep. 20, 356–369 (2017).

44. Rebane, A. A. et al. Liquid–liquid phase separation of the Golgi matrix protein GM130. FEBS Lett. 594, 1132–1144 (2020).

45. Chang, C.-C. et al. Ran pathway-independent regulation of mitotic Golgi disassembly by Importin-α. Nat. Commun. 10, 4307 (2019).

46. Ren, S., Sato, R., Hasegawa, K., Ohta, H. & Masuda, S. A Predicted Structure for the PixD–PixE Complex Determined by Homology Modeling, Docking Simulations, and a Mutagenesis Study. Biochemistry 52, 1272–1279 (2013).

47. Yuan, H. & Bauer, C. E. PixE promotes dark oligomerization of the BLUF photoreceptor PixD. Proc. Natl. Acad. Sci. 105, 11715–11719 (2008).

48. Dine, E., Gil, A. A., Uribe, G., Brangwynne, C. P. & Toettcher, J. E. Protein Phase Separation Provides Long-Term Memory of Transient Spatial Stimuli. Cell Syst. 6, 655–663.e5 (2018).

49. Chaurasia, S., Pieraccini, S., Gonda, R. D., Conti, S. & Sironi, M. Molecular insights into the stabilization of protein–protein interactions with small molecule: The FKBP12–rapamycin–FRB case study. Chem. Phys. Lett. 587, 68–74 (2013).

50. Banaszynski, L. A., Liu, C. W. & Wandless, T. J. Characterization of the FKBP·Rapamycin·FRB Ternary Complex. J. Am. Chem. Soc. 127, 4715–4721 (2005).

51. Lehotzky, A. et al. Dynamic targeting of microtubules by TPPP/p25 affects cell survival. J. Cell Sci. 117, 6249–6259 (2004).

52. Efimov, A. et al. Asymmetric CLASP-Dependent Nucleation of Noncentrosomal Microtubules at the trans-Golgi Network. Dev. Cell 12, 917–930 (2007).

53. Zhang, J., Mehta, S. & Zhang, J. Protocol for reading and imaging live-cell PKA activity using ExRai-AKAR2. STAR Protoc. 3, 101071 (2022).

54. Zhang, Q. et al. Visualizing Dynamics of Cell Signaling In Vivo with a Phase Separation-Based Kinase Reporter. Mol. Cell 69, 334–346.e4 (2018).

55. Massengill, C. I., Day-Cooney, J., Mao, T. & Zhong, H. Genetically encoded sensors towards imaging cAMP and PKA activity in vivo. J. Neurosci. Methods 362, 109298 (2021).

56. Hong, S. K., Kim, K.-H., Song, E. J. & Kim, E. E. Structural Basis for the Interaction between the IUS-SPRY Domain of RanBPM and DDX-4 in Germ Cell Development. J. Mol. Biol. 428, 4330–4344 (2016).

57. Castrillon, D. H., Quade, B. J., Wang, T. Y., Quigley, C. & Crum, C. P. The human VASA gene is specifically expressed in the germ cell lineage. Proc. Natl. Acad. Sci. 97, 9585–9590 (2000).

58. Zhu, F. et al. Deficiency of TPPP2, a factor linked to oligoasthenozoospermia, causes subfertility in male mice. J. Cell Mol. Med. 23, 2583–2594 (2019).

59. Coyle, S. M. Ciliate behavior: blueprints for dynamic cell biology and microscale robotics. Mol. Biol. Cell 31, 2415–2420 (2020).

60. Coyle, S. M., Flaum, E. M., Li, H., Krishnamurthy, D. & Prakash, M. Coupled Active Systems Encode an Emergent Hunting Behavior in the Unicellular Predator Lacrymaria olor. Curr. Biol. 29, 3838–3850.e3 (2019).

61. Teixidó-Travesa, N., Roig, J. & Lüders, J. The where, when and how of microtubule nucleation – one ring to rule them all. J. Cell Sci. 125, 4445–4456 (2012).

62. Bornens, M. The Centrosome in Cells and Organisms. Science 335, 422–426 (2012).

63. Rios, R. M. The centrosome–Golgi apparatus nexus. Philos. Trans. R. Soc. 369, 20130462 (2014).

64. Banani, S. F. et al. Compositional Control of Phase-Separated Cellular Bodies. Cell 166, 651–663 (2016).

65. Haque, F. et al. Cytoskeletal regulation of a transcription factor by DNA mimicry via coiled-coil interactions. Nat. Cell Biol. 24, 1088–1098 (2022).

66. Park, H. et al. Optogenetic protein clustering through fluorescent protein tagging and extension of CRY2. Nat. Commun. 8, 30 (2017).

67. Shih, P.-Y., Lee, S.-P., Chen, Y.-K. & Hsueh, Y.-P. Cortactin-binding protein 2 increases microtubule stability and regulates dendritic arborization. J. Cell Sci. 127, 3521–3534 (2014).

68. Zhou, H. et al. CCDC74A/B are K-fiber crosslinkers required for chromosomal alignment. BMC Biol. 17, 73 (2019).

