## Supplemental Materials for "Regulatable assembly of synthetic microtubule architectures using engineered MAP-IDR condensates"

Contains:
Supplementary Figures and Figure Legends 1-9
Supplementary Video Legends 1-8

Supplementary Figures and Figure Legends.

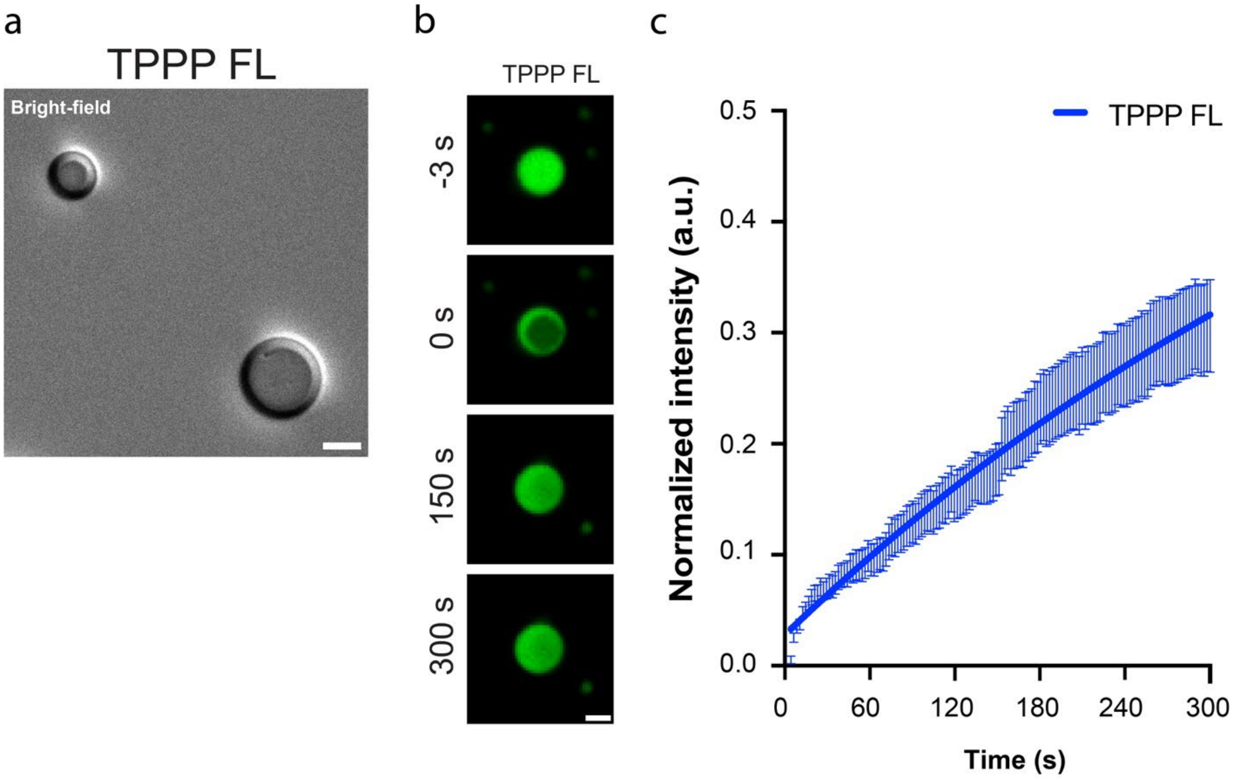


Supplementary Figure 1. **TPPP phase separates into a liquid-like condensate. a** Bright-field image of untagged TPPP protein (DIC) at the concentration of 20 μM with 20 mM HEPES, 50 mM NaCl, 3 mM DTT, and 12% dextran. Scale bars, 10 μm. **b** Representative images showing the liquid drops of GFP-TPPP, during FRAP (before bleaching, -3s; at bleaching, 0s; after bleaching, 150s and 300s). Scale bars, 2 μm. **c** FRAP of GFP-TPPP drops acquired by confocal microscopy. The dynamic recovery curves were determined by fitting to a One-phase association equation. (mean ± SEM, TPPP, n = 22) (see **Supplementary Video S1**).

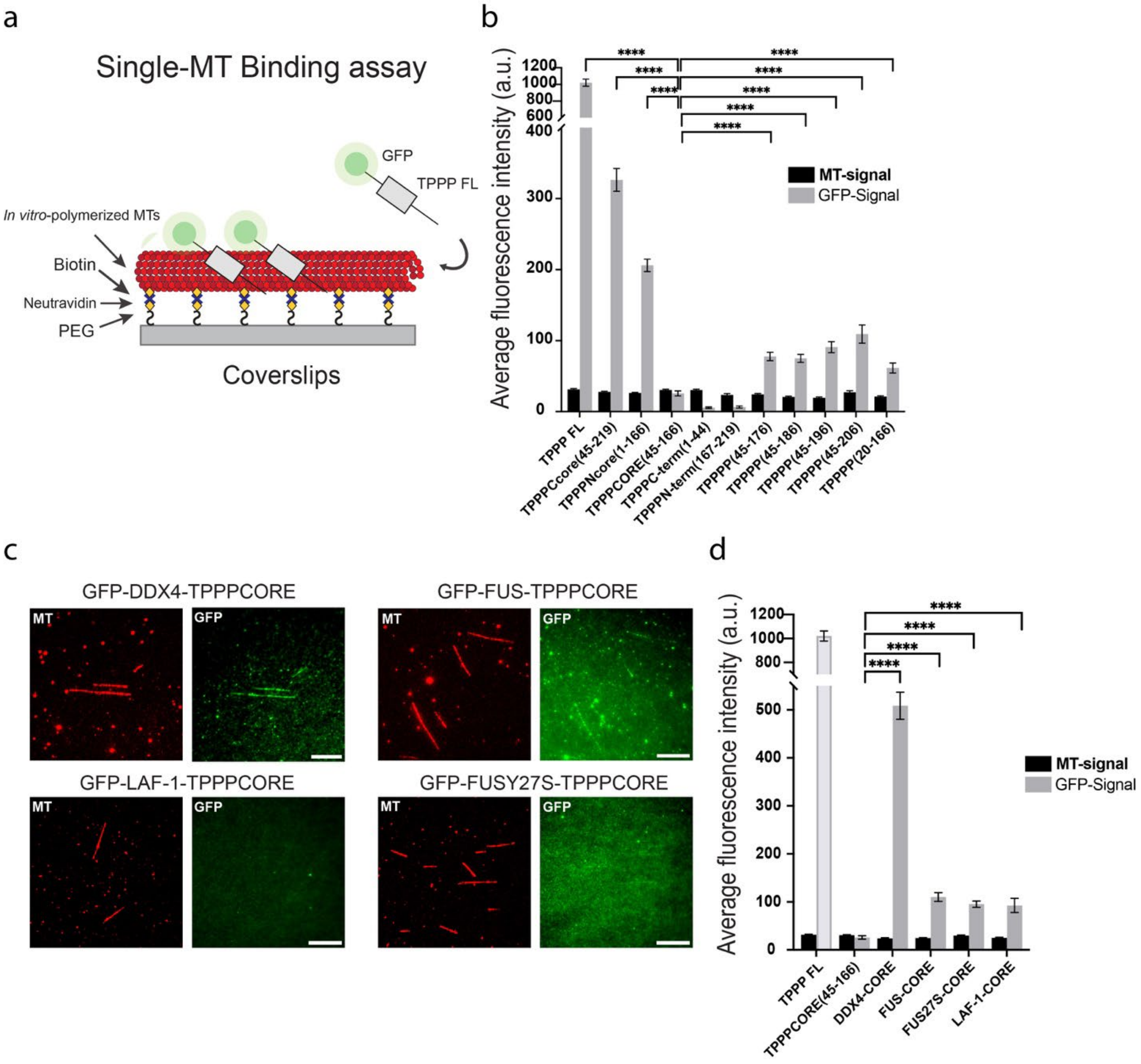


Supplementary Figure 2. **Single-MT binding assay for native TPPP variants and synMAP TPPP-IDR-chimeras. a** Schematic for single microtubule bundling assay. In vitro polymerized Alexa-594 (red), biotin-labeled, and GMPCPP-stabilized microtubules were immobilized on NeutrAvidin-coated biotinylated coverslip. After washing out extra microtubules, GFP-TPPP variants or GFP-TPPP-Chimeras (green) were added to the chamber in the concentration of 600 nM for 5 min. Followed by 1xBRB80 washing steps, the chamber was filled with the solution, sealed, and then imaged using TIRF microscopy. **b** Analysis of TPPP variants (GFP) and microtubule (Alexa-594) fluorescence signals. Mean fluorescence signals under the different conditions were measured and plotted. ( TPPP FL, n = 104; TPPP Ccore(45-219), n = 117; TPPP Core (1-166), n = 164; TPPP CORE(45-166), n = 52; TPPP(1-44), n = 31; TPPP(167-219), n = 47; TPPP(45-206), n = 47; TPPP(45-196), n = 46; TPPP(45-186), n = 77; TPPP(45-176), n = 84; TPPP(20-166), n = 68 microtubules for each conditions) SD was determined from data pooled. Two-tailed Student t test; statistical differences: ****, $P < 0.0001$; n.s., not significant. **c** GMPCPP-stabilized microtubules (Alexa-594 and biotin-labeled), immobilized on a glass surface, were incubated with TPPP Chimeras; GFP-DDX4-CORE, GFP-FUS-CORE, GFP-FUSY27S-CORE, and GFP-LAF-1-CORE, respectively. Scale bars, 10 µm. **d** Analysis of TPPP Chimeras (GFP) and microtubule (Alexa-594) fluorescence signals. Mean fluorescence signals under the different conditions were measured and plotted. (DDX4-CORE, n = 56; FUS-CORE, n = 76; FUSY27S-CORE, n = 52; LAF-1-CORE, n = 38 microtubules for each conditions) SD was determined from data pooled. Two-tailed Student t test; statistical differences: ****, P < 0.0001; n.s., not significant.

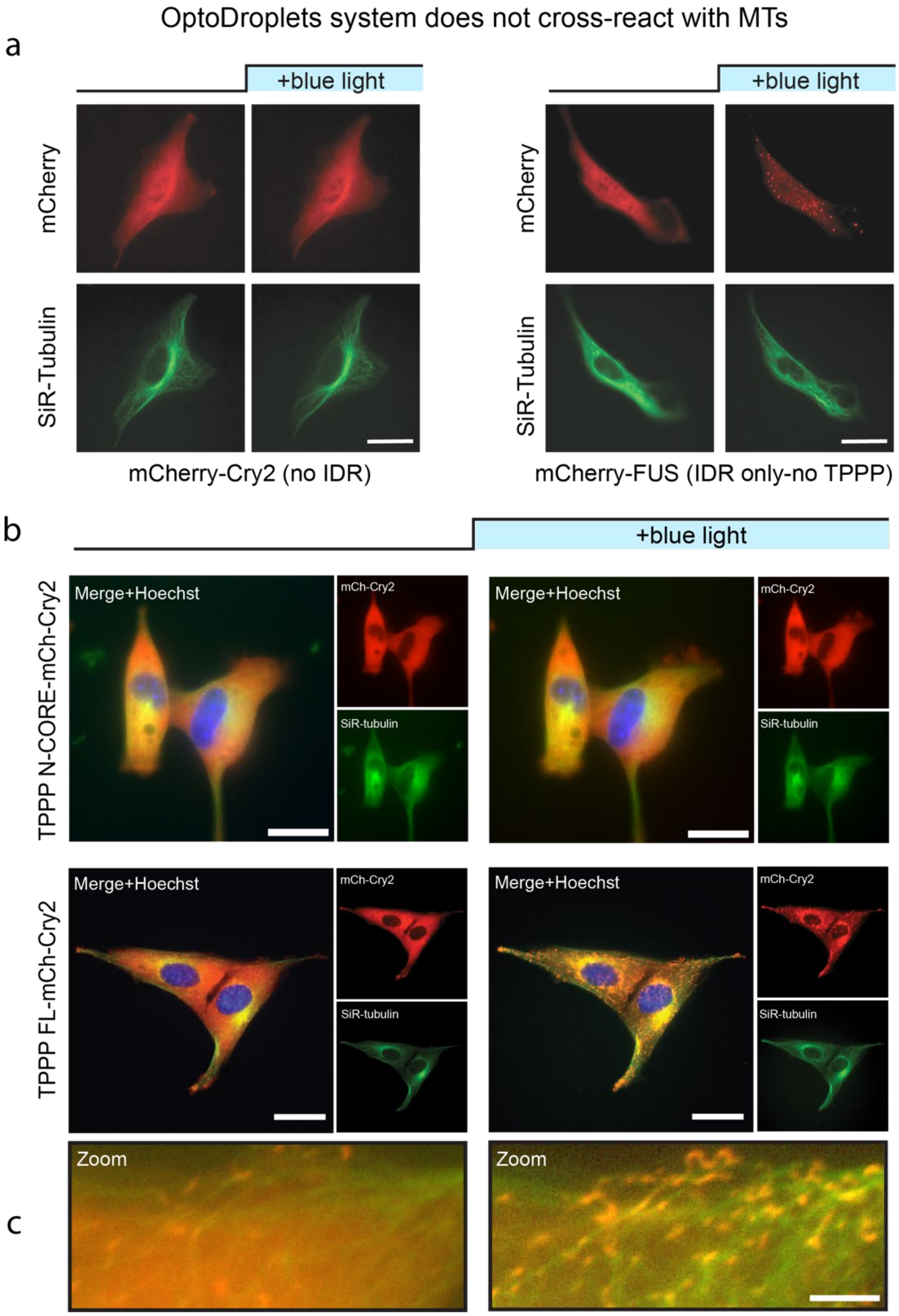


Supplementary Figure 3. **Optical clustering of TPPP stimulates microtubule (MT) association but not MT bundling.** Representative images of pre- and post-blue light activation 5 min of **a** (left) Cry2, **a** (right) OptoFUS, **b** OptoTPPPNcore and OptoTPPP FL in NIH3T3 cells. Scale bars, 20 µm. **c** Higher-magnification images of OptoTPPP FL a. Scale bar, 5 µm. (see Supplementary Video S2).

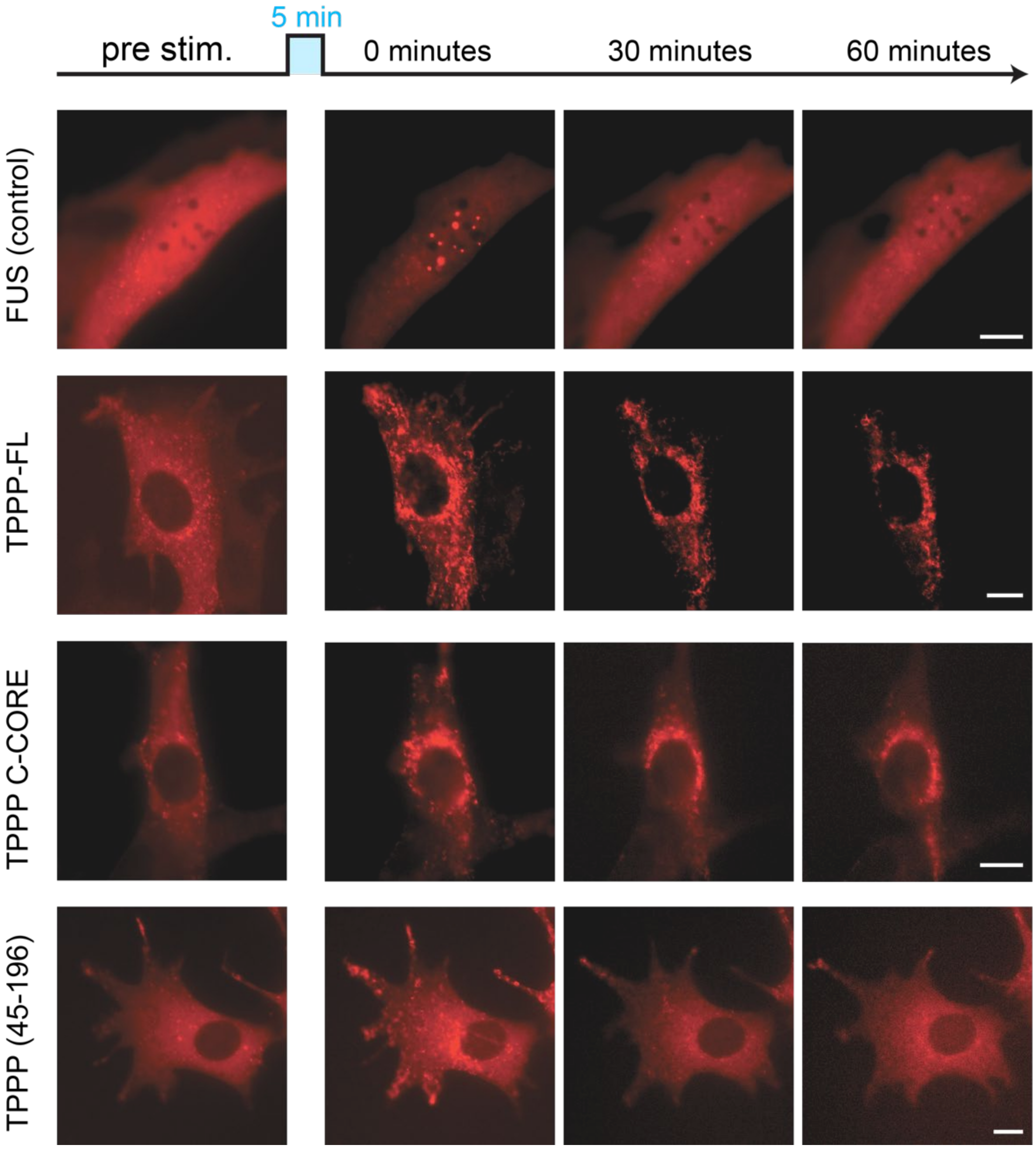


Supplementary Figure 4. **Disassembly of optically-induced clusters formed by different TPPP variants.** Representative images of OptoFUS and OptoTPPP cells under cluster disassembly assay. Blue box indicated as blue light activation for 5 min and then incubated in the absence of blue light for 60 min. Cell images are captured during (pre-activation, after light stimulation; 0s, 30 min and 60 min). Scale bar, 10 μm. Related to **Figure 3D.**

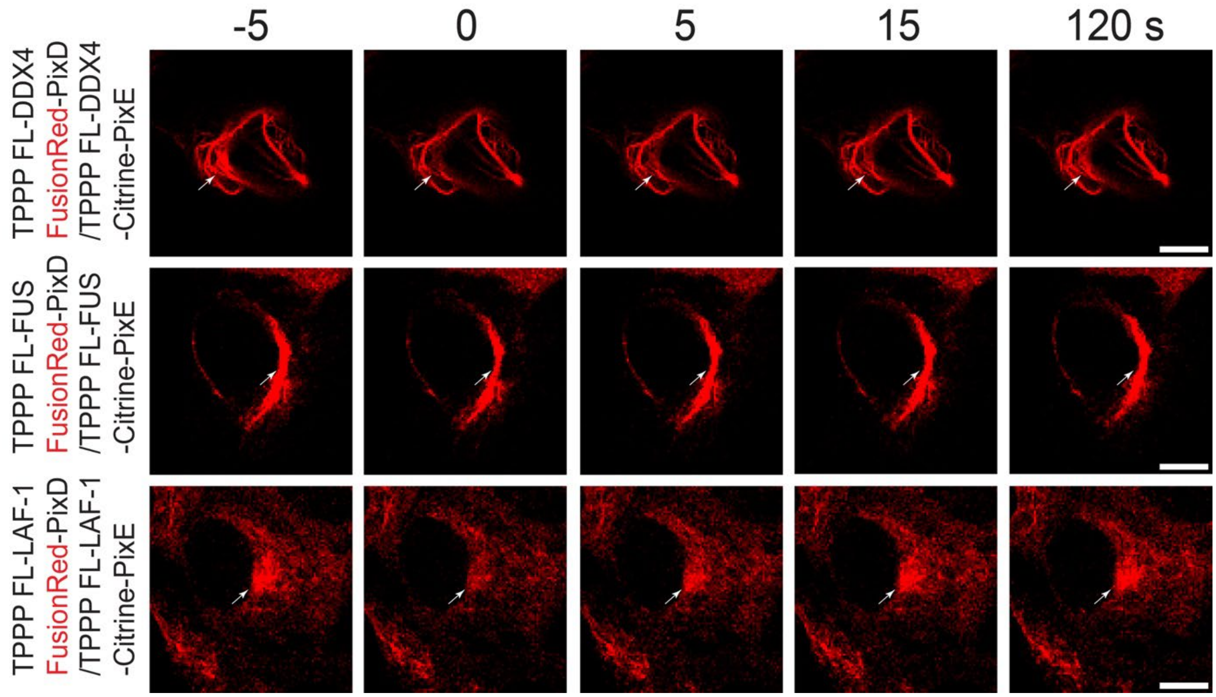


Supplementary Figure 5. **FRAP analysis of synMAP TPPP-IDR-PixDE bundled MTs in NIH3T3 cells.** Representative sequential confocal image of different TPPP FL-IDR-PixDE bundled MTs (FusionRed, red) in NIH3T3 cells during FRAP (before bleaching, -5s; at bleaching, 0s; after bleaching, 5s, 15s and 120s). Scale bars, 10 μm. Related to **Figure 4H.**

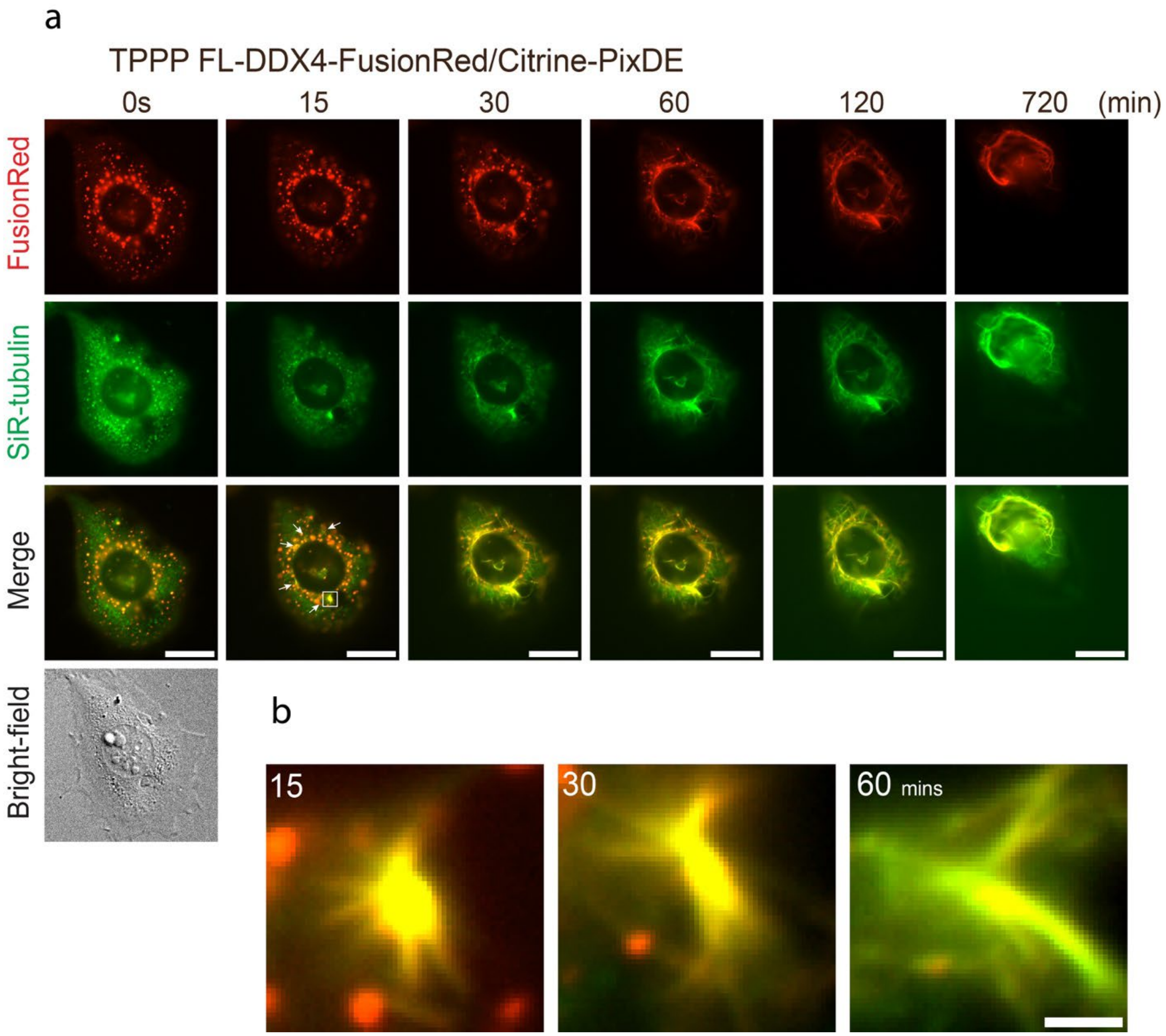


Supplementary Figure 6. **synMAP TPPP-IDR condensates can nucleate MTs in mammalian cells. a,b** Video frames illustrating MT formation at TPPP phase-condensed hub(DDX4-PixDE) in NIH3T3 cell. After Nocodazole washout, the TPPP-DDX4-PixDE droplets (FusionRed,red and SiR-tubulin, green) start to nucleate and bundle up MTs. Time after nocodazole removal is shown. Scale bar, 20 μm. **b** Higher-magnification images of TPPP phase-condensed hub MT-nucleation, Scale bar, 2 μm. (see **Supplementary Video S6**).

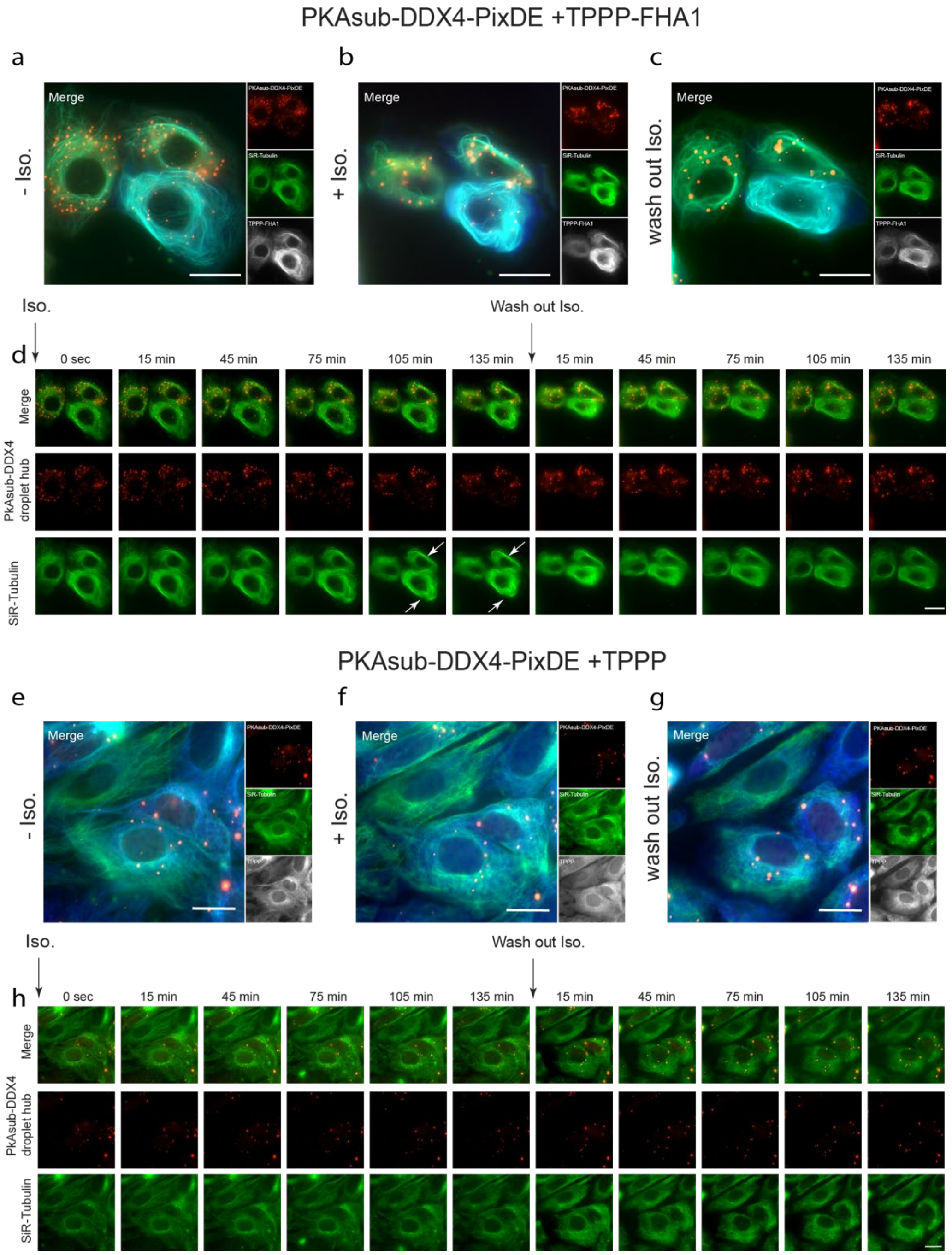


Supplementary Figure 7. **Images from timecourse of synMAP circuit behavior in which MT architecture is regulated by Protein Kinase A (PKA) signaling activity**. NIH3T3 cells co-transfected with PKAsub-DDX4-PixDE and TPPP-FHA1 or TPPP were treated with 20 μM isoprenaline (Iso) for 135 minutes and then washed out. **a-c, e-g** Epifluorescent images of PKAsub-DDX4-PixDE (red) and TPPP-FHA1 or TPPP MTs (tagBFP, white color in image) before, after and wash out isoprenaline treatment. SiR-tubulin (green) is a marker for MT lattices. Scale bar, 20 μm. **d,h** Representative images of a time series showing the recruitment of TPPP-FHA1 or TPPP (control) to PKAsub-DDX4-PixDE (FusionRed, red) droplets induced by isoprenaline. Microtubule lattices were stained with SiR-tubulin (green). Scale bar, 20 μm.

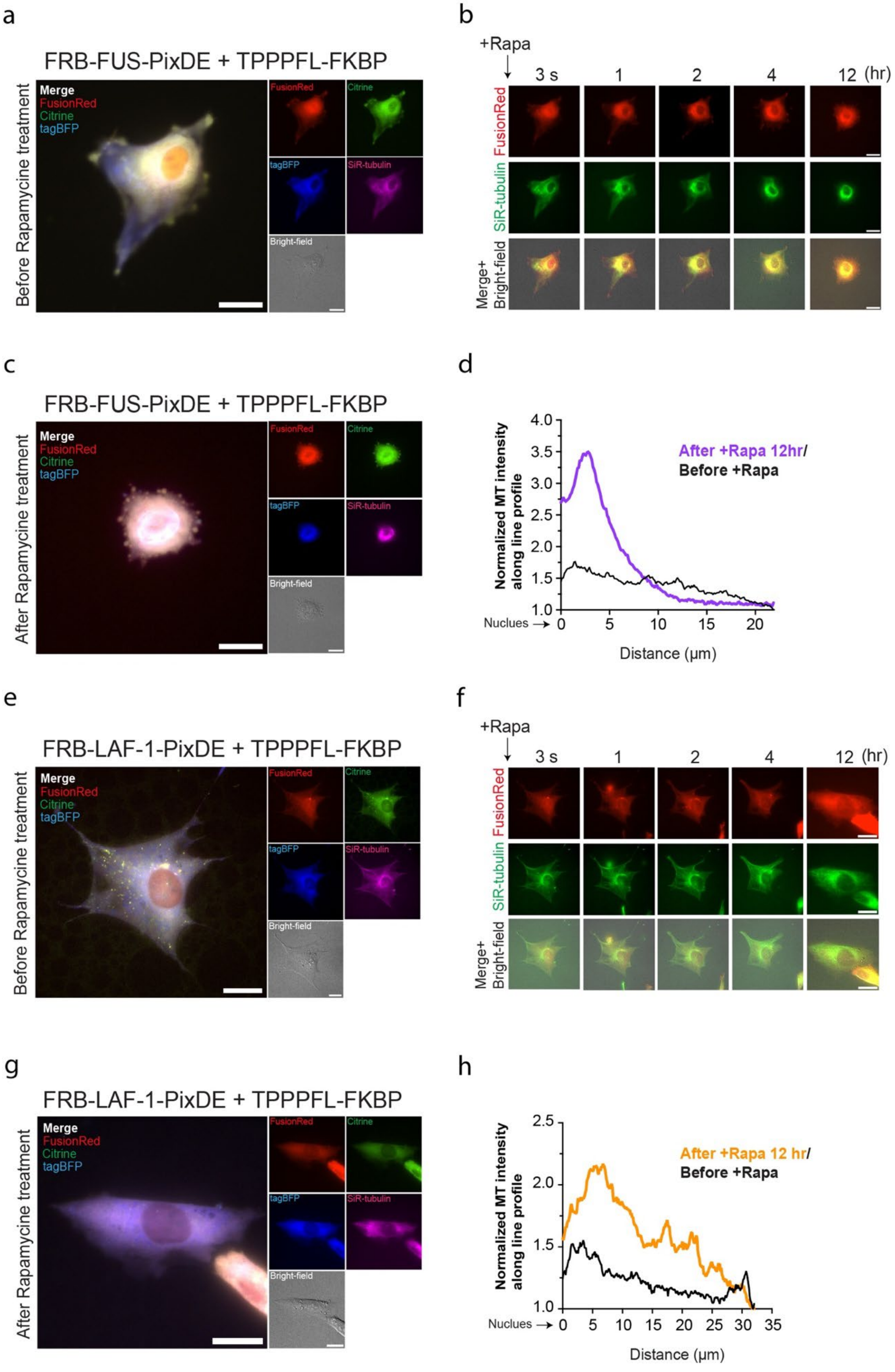
a
FRB-FUS-PixDE + TPPPFL-FKBP
Before Rapamycine treatment
Merge
FusionRed
Citrine
tagBFP
SiR-tubulin
Bright-field
b
+Rapa
3 s
1
2
4
12 (hr)
FusionRed
SiR-tubulin
Merge+
Bright-field
c
FRB-FUS-PixDE + TPPPFL-FKBP
After Rapamycine treatment
d
Normalized MT intensity along line profile
After +Rapa 12hr/
Before +Rapa
Nuclues →
Distance (μm)
e
FRB-LAF-1-PixDE + TPPPFL-FKBP
Before Rapamycine treatment
f
g
FRB-LAF-1-PixDE + TPPPFL-FKBP
After Rapamycine treatment
h
After +Rapa 12 hr/
Before +Rapa

Supplementary Figure 8. **Images and single-cell quantification of behavior of inducible synMAP circuits that vary in the identity of the condensate-forming component.** NIH3T3 cells co-transfected with FRB-FUS-PixDE or FRB-LAF-1-PixDE and TPPP-FKBP were treated with 20 µM rapamycin (Rapa). Before and after rapamycin treatment. **a,c,e,g** Epifluorescent images of FRB-FUS-PixDE (FusionRed, red) (Citrine, green) or FRB-LAF-1-PixDE (FusionRed, red) (Citrine, green), TPPP-coated MTs (tagBFP, blue), and DIC. SiR-tubulin (purple) is a marker for MT lattices. Scale bar, 20 µm. **b,f** Representative image of a time series of rapamycin-induced TPPP into either FRB-FUS-PixDE (FusionRed, red) or FRB-LAF-1-PixDE drops (FusionRed, red) condensed hubs. Microtubule lattices were stained with SiR-tubulin (green). Scale bar, 20 µm. **d,h** Quantification of the microtubule fluorescence signal (SiR-tubulin) before and after 12 hr rapamycin treatment distribution in NIH3T3 cells co-transfected with FRB-FUS-PixDE or FRB-LAF-1-PixDE and TPPP-FKBP. Distance from the center of the Nucleus is shown on the horizontal axis, and normalized fluorescence intensity is on the vertical axis.

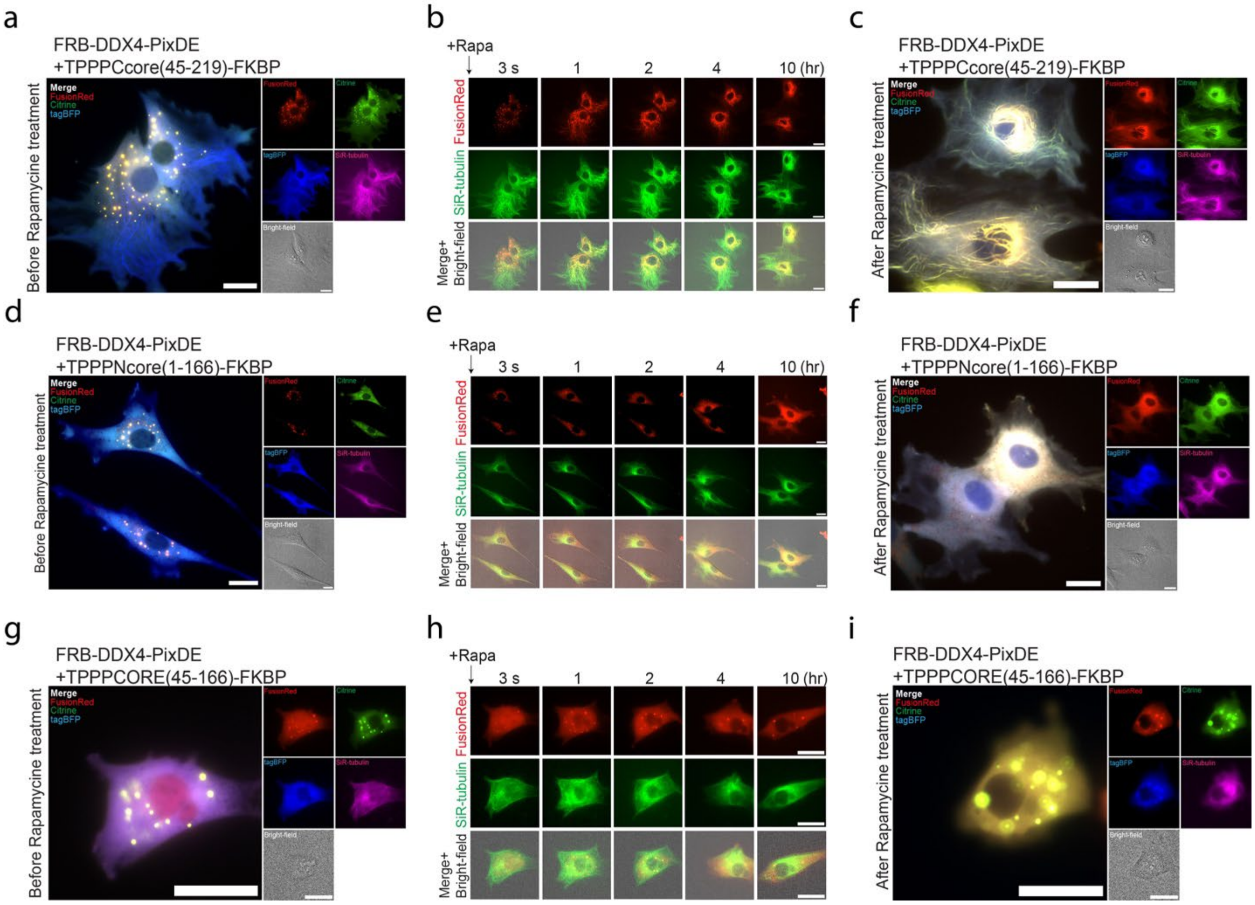


Supplementary Figure 9. **Images and single-cell quantification of behavior of inducible synMAP circuits that vary in the identity of the microtubule-interacting component.** NIH3T3 cells co-transfected with FRB-DDX4-PixDE and TPPPCore(45-219) or TPPPNcore(1-166) or TPPPCORE(45-166) were treated with 20 µM rapamycin (Rapa). Before and after rapamycin treatment. **a,c,d,f,g,i** Epifluorescent images of FRB-DDX4-PixDE (FusionRed, red) (Citrine, green), TPPP variant with tagBFP-FKBP (tagBFP, blue), and DIC. SiR-tubulin (purple) is a marker for MT lattices. Scale bar, 20 µm. **b,e,h** Before and after 10 hr rapamycin treatment in NIH3T3. Representative images of a time series of rapamycin-induced either TPPPCore or TPPPNcore or TPPPCORE into FRB-DDX4-PixDE droplet hubs (FusionRed, red). Microtubules were stained with SiR-tubulin (green). Scale bar, 20 µm.

**Supplementary Video S1** Time-lapse of the Fluorescent Recovery After Photo-bleaching (FRAP) of GFP-TPPP FL drops. TPPP drops were formed with 20 µM GFP-TPPP FL, 20 mM HEPES, 50 mM NaCl, 3 mM DTT, 12% dextran, pH 7.4.

**Supplementary Video S2.** Cluster formation of TPPP FL-mCh-Cry2WT in NIH3T3 cell. OptoTPPP(red) forms clusters, condensing on MTs after blue-light activation. mCh (red) and SiR-tubulin (green).

**Supplementary Video S3.** Cluster formation assay of TPPPCORE-mCh-Cry2WT in NIH3T3 cell. Removal of IDRs of TPPP led to inhibition of the cluster formation in the living cell. mCh (red) and SiR-tubulin (green).

**Supplementary Video S4.** Rapid microtubule bundling formation by recruiting TPPP into a phase-condensed hub (DDX4-PixDE), FusionRed (red) and SiR-tubulin (green). Related to **Figure 5A-D.**

**Supplementary Video S5.** The TPPP-induced MT bundles could be maintained and passed into two daughter cells. (TPPP-FKBP/FRB-DDX4-PixDE) FusionRed (red) and SiR-tubulin (green).

**Supplementary Video S6.** Microtubule nucleation and bundling from the TPPP phase-condensed hub (DDX4-PixDE) in NIH3T3 cells after Nocodazole Washout. TPPP-DDX4-FusionRed/Citrine-PixD/E(red) and SiR-tubulin (green). Related to **Figure 5E,F and Supplementary Figure 6.**

**Supplementary Video S7.** PKA signaling control of microtubule bundling formation in the NIH3T3 cells. (TPPP-FHA1/PKAsub-DDX4-PixDE) FusionRed (red) and SiR-tubulin (green). Related to **Figure 5G-H and Supplementary Figure 7.**

**Supplementary Video S8.** Recruiting TPPPCORE to the intense phase-condensed hubs (DDX4-PixDE) could partially restore the MTs-bundling activity in NIH3T3 cells. (TPPPCORE-FKBP/FRB-DDX4-PixDE) FusionRed (red) and SiR-tubulin (green).
